# Whole genome sequencing of newly emerged fungal pathogen *Aspergillus lentulus* and its Azole resistance gene prediction

**DOI:** 10.1101/2023.12.28.573576

**Authors:** Xiaodong Wang, Aikedai Yusufu, Hadiliya Hasimu, Paride Abliz

## Abstract

*Aspergillus lentulus* is an important newly recorded species in the *A. fumigatus* complex and its resistance to azole drugs and the high mortality rate of infected individuals have emerged as problems. Comprehensive understanding of the *A. lentulus* is limited due to lack of genome-wide fine mapping data. The aim of this study was to investigate the *A. lentulus* signature at the molecular level, analyze the genome-wide profile of this strain and predict its possible genes that execute azole resistance. In this study, a whole genome sequencing of a clinically isolated *A. lentulus* strain (named *A. lentulus* PWCAL1) was studied by PacBio Sequel sequencing platform. Azole resistance genes were predicted based on whole-genome sequencing data analysis, gene function annotation, comparative genomics analysis, and blastp alignment using the Mycology Antifungal Resistance Database (MARDy) to comprehensively understanding the genome-wide features, pathogenicity, and resistance mechanisms of *A. lentulus*. The results of whole genome sequencing demonstrated that the total length of *A. lentulus* PWCAL1 genome was 31255105 bp, the GC content was 49.24%, and 6883 coding genes were predicted. A total of 4565, 1824 and 6405 genes were annotated in GO, KOG and KEGG databases, respectively. In the PHI and DFVF databases, 949 and 259 interacting virulence factors were identified, respectively, with the main virulence factor-Mutant virulence phenotype was enriched in reduced virulence. Comparative genomics analysis showed that there were 5456 consensus Core Genes in this strain and four closely related strains of *A. fumigatus* complex, which were mainly involved in Human Diseases, Metabolism, Organismal Systems, etc. Among the three aligned *A. lentulus* strains, the number of unique genes of this bacterium was the highest with number of 171, and these genes were mainly associated with Carbohydrate metabolism, Cell growth and death. Resistance gene prediction demonstrated that the A5653 gene of this bacterium had F46Y/N248T double point mutations on the CYP51A gene, but no tandem repeat TR mutations in the promoter region were detected. Further more, twelve genes belonging to the fungal multidrug resistance ABC transporters were identified based on the complete genome sequence and phylogenetic analysis of A. lentulus, which belonged to the ALDp subfamily, the PDR subfamily (AtrB, CDR1, and CDR2), and the MDR subfamily (MDR1), respectively, and there were four genes that are annotated to the MFS multidrug transporter. Further phylogenetic tree classification of the ABC transporter subfamilies predicted in the nine selected *A. fumigatus* complex strains showed that these putative ABC proteins were divided into two main clusters, which belonged to PDR (CDR1, CDR2, AtrB, and AtrF) and MDR subfamilies (MDR1, MDR2, and MDR3). The distribution of these ABC proteins varies among different species of the *A. fumigatus* complex. The main result obtained from this study for the whole genome of *A. lentulus* provide new insights into better understanding the biological characteristics, pathogenicity and resistance mechanisms of this bacterium. In this study, two resistance mechanisms which include CYP51A gene mutation and multidrug-resistant ABC transporter were predicted in a single isolate. Based on the predicted CYP51A-F46Y/N248T site mutation combination, we speculate that the CYP51A gene of *A. lentulus* may be partially responsible for azole resistance. Based on the predicted ABC transporter family genes, we hypothesize that resistance to multiple azoles in *A. lentulus* is mediated, at least in part, by these ABC transporters with resistance.

## 1. Introduction

With the development of molecular biology, *A. fumigatus* identified by traditional morphological methods was found to be an *A. fumigatus* complex by using molecular identification methods, and cryptic species in these complexes can cause severe invasive fungal infections [Sindhu D et al.2019;Pinto E et al.2018]. Currently molecularly identified *A. fumigatus complex* species are *A. fumigatus, A. lentulus*, and *A. thermomutatus* and so on [Alcazar-Fuoli L et al.2008;Lamoth F.2016]. Among them, *A. lentulus* is a new species in the *A. fumigatus* complex first named by molecular identification in 2005 by Balajee et al., and sibling species with *A. fumigatus* [Balajee SA et al.2005]. In recent years, increasing cases of *A. lentulus* infection have been reported [Ahmed J et al.2022;Nematollahi S et al.2021;Datta K et al.2013], and recently, a study in China reported that six of 580 filamentous fungi were molecularly identified as *A. lentulus* [Yu SY et al.2020]. However, worldwide epidemiological data are limited.

*A. lentulus* is morphologically indistinguishable from *A. fumigatus*, which made difficulty in clinical diagnosis. Unlike *A. fumigatus*, sporulation of *A. lentulus* is slow and small and does not grow at 48℃. The only draft genome of *A. lentulus* IFM54703T currently shows that the genome size of this bacterium is 30,956,128 bp and G + C content is 49.45% [Kusuya Y et al.2016]. But the complete genome-wide fine map of *A. lentulus* has not been reported.

Many clinical reports have shown that *A. lentulus* infection is mainly associated with invasive pulmonary aspergillosis. Infected patients are mainly organ transplant recipients, immunocompromised individuals, and these infections tend to cause fatal outcomes[Bastos VR et al.2015;Alhambra A et al;2008;Yagi K et al.2019;Dos Santos RAC et al.2020]. The main concern in clinical treatment is that *A. lentulus* has reduced sensitivity or even resistance to multiple antifungal drugs (azoles, polyenes, and echinocandins), especially azoles, and treatment failure leads to a high mortality rate[Balajee SA et al.2005]. In China, Yu et al. reported that the MIC values of in vitro antifungal susceptibility testing of 6 clinically isolated *A. lentulus* demonstrated that amphotericin B was 4-8 μg/ml, itraconazole was 2 μg/ml, voriconazole was 4-16 μg/ml, and posaconazole was 0.5-1 μg/ml [Yu SY et al.2020]. It is therefore essential to identify the exact mechanism of resistance in *A. lentulus*, particularly azole resistance.

The specific resistance mechanism of *A. lentulus* to azoles is still unknown. Studies have confirmed that the mechanism of azole resistance in *Aspergillus* is mainly a combination of single point mutations in the CYP51A gene and tandem repeat (TR) mutations in the promoter region. In addition, different combinations of five Cyp51A mutations (F46Y, M172V, N248T, D255E, and E427K) have been reported worldwide in approximately 10% of all *A. fumigatus* isolates tested [Zarrin M et al.2018;Gonzalez-Jimenez I et al.2021;Nargesi S et al.2022].

Although the most prevalent mechanism of azole resistance is alteration of CYP51A, many resistant isolates without mutations or alterations in CYP51A or its promoter have been reported[Bowyer P et al.2011].

Another important mechanism that related to the azole resistance in *Aspergillus* is overexpression of the efflux system, and the associated genes are mainly ATP-binding cassette transporter (ABC) and the major facilitator superfamily (MFS) transporter, and rapid active drug extrusion represents cells from one of the major mechanisms of multidrug resistance [Cannon RD et al.2009]. Studies have shown that the currently annonated *A. fumigatus* genome includes 45 ABC family transporters and 275 MFS family transporters [Meneau I et al.2016]. Currently known transporters that are related to the azole resistance in *Aspergillus* includes mdr1, mdr2, mdr3, mdr4, AtrF, AtrB, abcD, abcE, and cdr1B. *A. lentulus* is definitely related to *A. fumigatus*, but regarding the mechanism of azole resistance mediated by the efflux pump of *A. lentulus* is still unknown. Based on the efflux transporter similarity with *C. Alaicans* ABC-transporters(cdr1,cdr2 and cdr4) and MFS transporters(mdr1) recent evidence suggested a relationship between the expression of cdr1B and itraconazole resistance in non-Cyp51A-mediated *A. fumigatus* isolates[Fraczek MG et al.2013].

In this study, we present the first genome-wide sequencing study of clinically isolated *A. lentulus* and accurately interpreted the sequence information. Clinically isolated *A. lentulus* in Xinjiang which is characterized with CYP51A-F46Y/N248T mutation combination was identified by Gene annotation and comparative analysis, and 12 azole resistance-associated ABC transporters were predicted, the ABC transporters predominantly belong to the PDR, MDR, and ALDp subfamilies. Phylogenetic tree comparative analysis have also demonstrated that there were 8 ABC transporters of *A. fumigatus* complex species which predominantly belong to PDR and MDR, and these efflux pump encoding genes may play a role in reducing azole susceptibility.

## 2. Materials and Methods

### 2.1. Fungal strain and DNA extraction

In this study, the experimental strain *A. lentulus* was isolated from sputum of a patient with chronic obstructive emphysema (COPD) and named *A. lentulus* PWCAL1, which had MIC values of 2 ug/mL for voriconazole and 4 ug/mL for amphotericin B. The strains were cultured in PDA medium at 37℃ for 7 days, and genomic DNA was extracted by SDS, detected by agarose gel electrophoresis, and quantified by Qubit ® 2.0 fluorescence quantifier (Thermo Scientific).

### 2.2. Whole Genome sequencing and functional annotation

Whole genome sequencing was conducted by PacBio Sequel platform. Seven databases were used to predict the gene function, which includes GO (Gene Ontology), KEGG (Kyoto Encyclopedia of Genes and Genomes), KOG (Clusters of Orthologous Groups), NR (Non-Redundant Protein Database), TCDB (Transporter Classification Database), P450, and Swiss-Prot. Genome-wide Blast searches (e-value less than 1e-5 and minimum alignment length percentage greater than 40%) were performed on the above seven databases. Secretory proteins were predicted by Signal P database. Meanwhile, secondary metabolic gene clusters were analyzed by anti-smash. Pathogenicity and resistance analyses were performed using PHI (Pathogen Host Interactions Database) and DFVF (database of fungalvirulence factors). Carbohydrate-active enzymes were predicted by CAZy (Carbohydrate-Active enZYmes Database). The sample genomes were visualized using Circos software.

### 2.3. Comparative Genomics Analysis

Cluster analysis of the protein sequences of the analyzed strains using CD-HIT software for core gene and specific gene analysis, and all non-redundant gene sets obtained by clustering were mapped to Pan Gene, extract the gene set shared by the 5 comparison strains in the clustering results as Core Gene, and the specific gene set in a single strain as Specific Gene. According to the distribution of each gene set in the genomes of five aligned strains, Pan Gene Wenckebach plots were drawn to demonstrate clustering between strains. Both core genes and specific genes were functionally annotated using KEGG and KOG databases

### 2.4. Screening and Analysis of Drug Resistance Genes

CYP51A resistance mutation analysis was performed using the Mycology Antifungal Resistance Database (MARDy) database for blastp alignment [Moazeni M et al.2020], and ABC transporter superfamily resistance genes were analyzed using TCDB database annotation. Establishment of Phylogenetic tree for ABC protein subfamily classification: First, protein sequences from nine strains were extracted and aligned with mafft, and then NJ evolutionary tree was constructed with treebest with bootstrap value of 1000.

### 2.5. Ethical Approval

The study was conducted in accordance with the Declaration of Helsinki. Ethical approval for the study was obtained by the Ethics Committee of the First Affiliated Hospital of Xinjiang Medical University. All patients were consent to be involved in this study.

## 3. Results and discussion

### 3.1. Basic gemonic characteristics of *A. lentulus*

After optimizing the genome assembly results of *A. lentulus* PWCAL1 strain and de novo Augustus prediction, 16 contigs were finally obtained, with a whole genome size of 31255105 bp and a GC content of 49.24%, which was basically consistent with the previous findings of 30956128 bp (accession number: BCLY01000001) and 31229376 bp (accession number: JAAAPU000000000) [Kusuya Y et al.2016;Dos Santos RAC et al.2020]. A total of 6883 coding genes were predicted (see Table 1-2) and their genome-wide circle plots are presented in Figure 1.The gene average length is 1334 bp (see Figure 2) In addition, the coding gene prediction results, tandem repeats, and non-coding RNA information of this bacterium are shown in Tables 2-5, respectively. However, no miRNAs were observed in *A. lentulus* PWCAL1 (see Table 5), and probably no fungal miRNA genes have yet been documented in the miRNA database.

**Figure 1.**
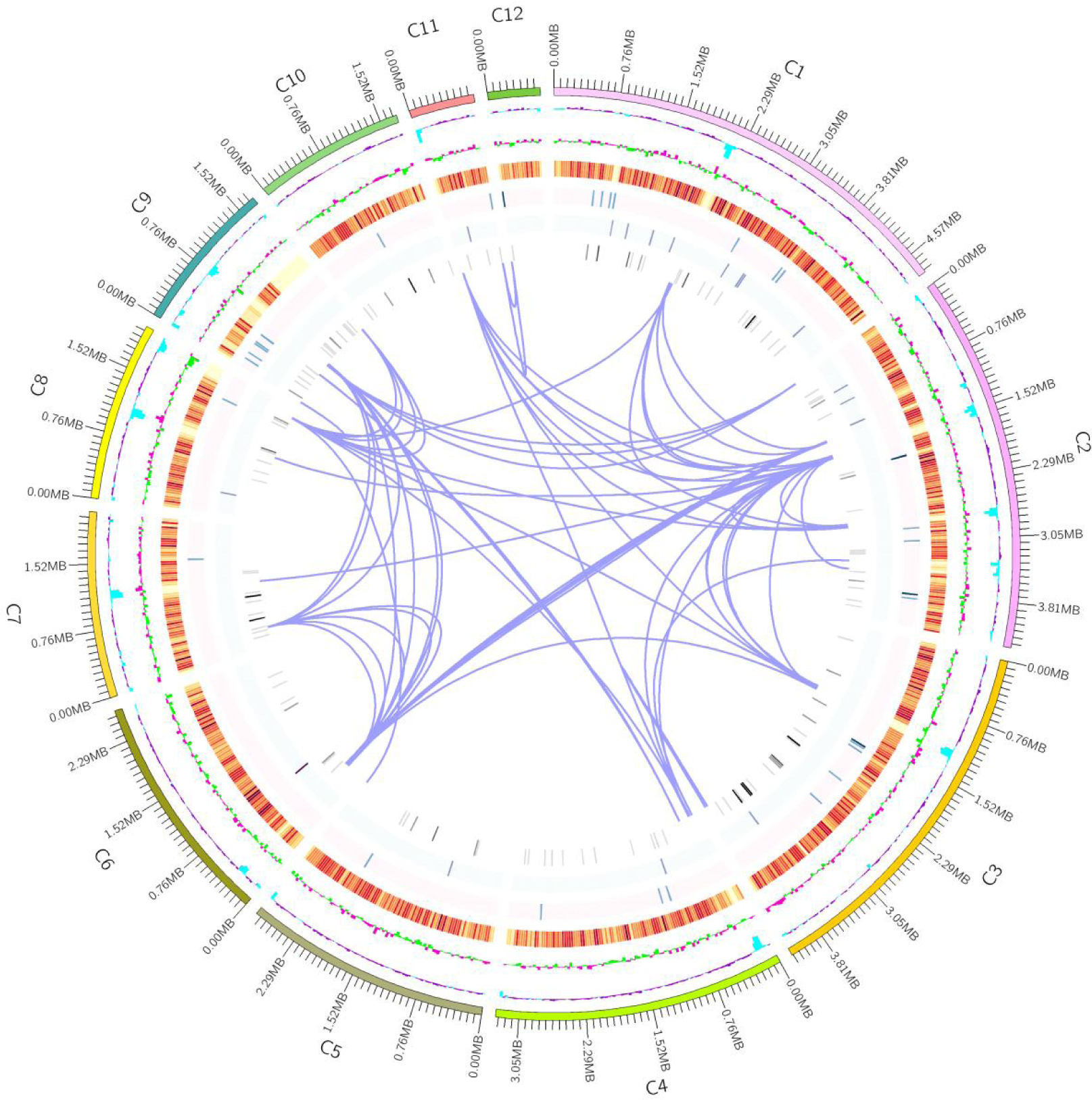
Genomic cycle diagram of the *A. lentulus*. Note: The outer circle represents the coordinates of the genome sequence position, from outside to inside, which sequentially represents GC content, GC skew, (gene density of coding genes, rRNA, snRNA and tRNA), and Gene Duplication. Genomic GC content: The GC content is calculated using a window (chromosome length/1000) bp and a step size (chromosome length/1000) bp. The sky blue portion inward indicates that the GC content in the region is lower than the average GC content of the entire genome, while the dark purple portion outward is on the contrary, as the peak goes higher, the greater the difference between the GC content and the average GC content. Genomic GC skew value: window (chromosome length/1000) bp, step size (chromosome length/1000) bp, The specific algorithm is G-C/G+C. The inward light green part indicates that the content of G in the region is lower than that of C, while the outward pink part indicates contrary result. Gene density: window (chromosome length/1000) bp, step length (chromosome length/1000) bp, and the proportion of genes in each window. Calculate the gene density of coding genes, rRNA snRNA tRNA, respectively. The darker the color, the greater the gene density within the window, and chromosome duplication.

**Figure 2.**
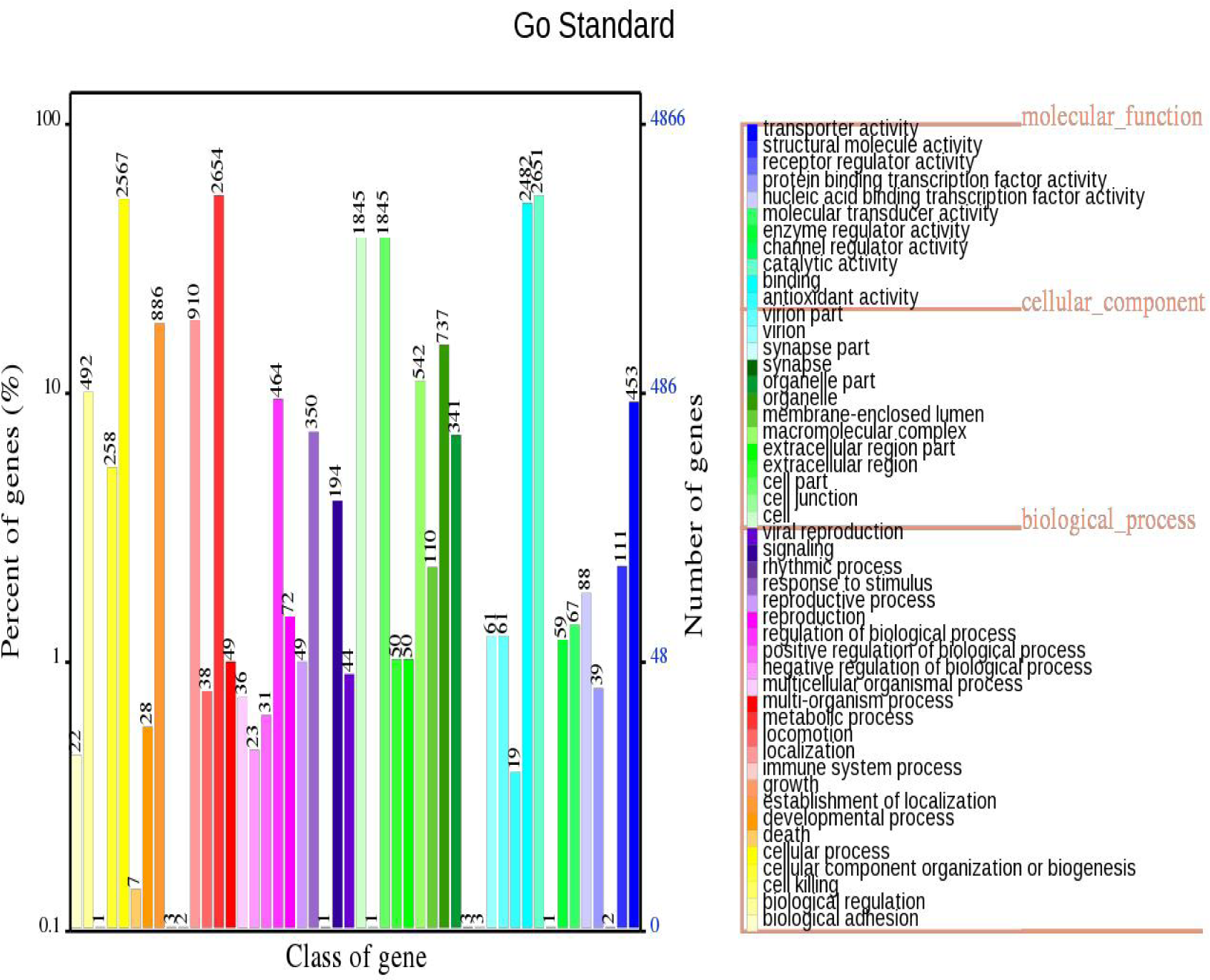
*A. lentulus* gene length distribution map

**Table 1.**
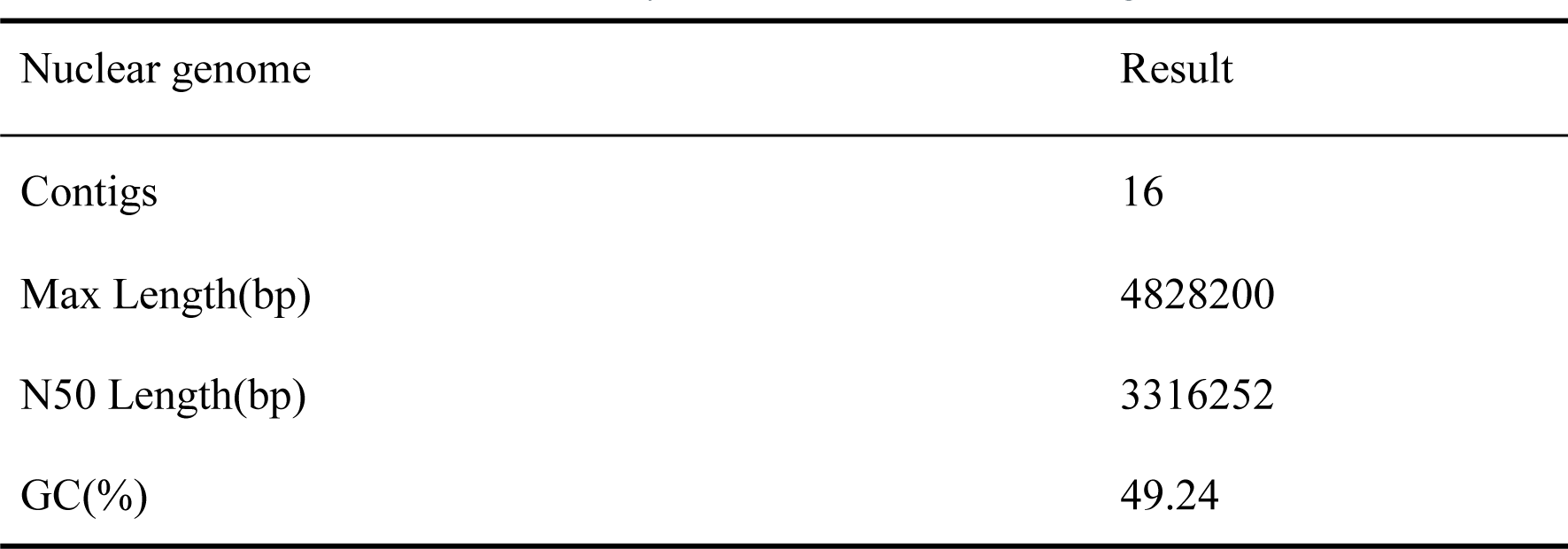
Final assembly results of the *A. lentulus* genome.

| Nuclear genome | Result |
| --- | --- |
| Contigs | 16 |
| Max Length(bp) | 4828200 |
| N50 Length(bp) | 3316252 |
| GC(%) | 49.24 |

**Table 2.** Prediction results of the *A. lentulus* coding genes.

| Nuclear genome | Result |
| --- | --- |
| Genome Size(bp) | 31255105 |
| Gene Number | 6883 |
| Gene Length(bp) | 9181152 |
| GC Content | 54.73 |
| % of Genome(Genes) | 29.37 |
| Gene Average Length | 1334 |
| Gene Internal Length | 22073953 |
| Gene Internal GC Content | 46.97 |
| % of Genome(internal) | 70.63 |

**Table 3.** Statistics of the *A. lentulus* scattered repeat sequence results.

| Type | Number | Total<br>Length(bp) | In Genome(%) | Average length(bp) |
| --- | --- | --- | --- | --- |
| LTR | 1107 | 279250 | 0.8935 | 253 |
| DNA | 422 | 85162 | 0.2725 | 206 |
| LINE | 346 | 38539 | 0.1233 | 116 |
| SINE | 117 | 8495 | 0.0272 | 74 |
| RC | 30 | 3461 | 0.0111 | 115 |
| Unknown | 7 | 491 | 0.0016 | 70 |
| Total | 2029 | 410045 | 13119 | 207 |

Note: LTR: Long Tandem Repeat; DNA: DNA Transposable; LINE:Long Interspersed Nuclear Element; SINE:Short Interspersed Nuclear Element; RC: Rolling Circle

**Table 4.**
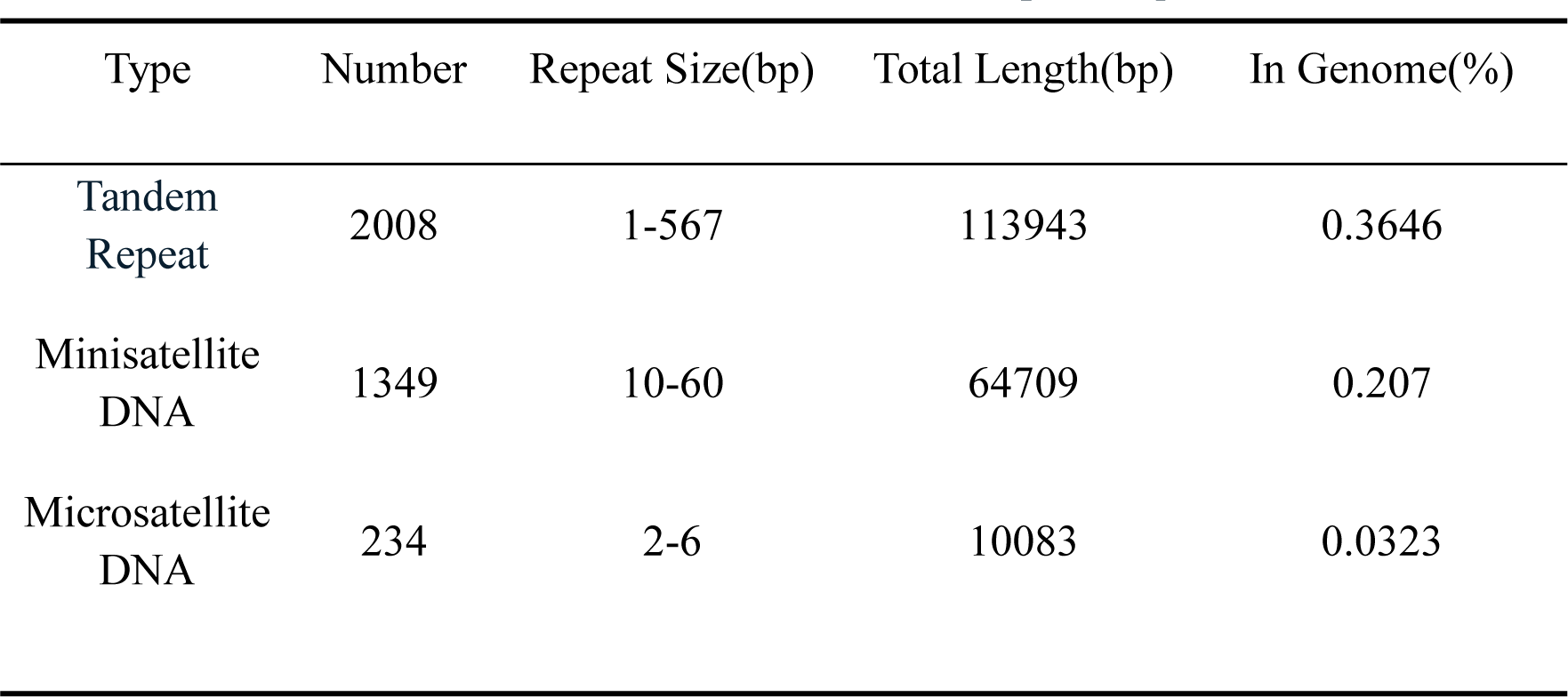
Statistics of the *A. lentulus* tandem repeat sequence results.

**Table 5.**
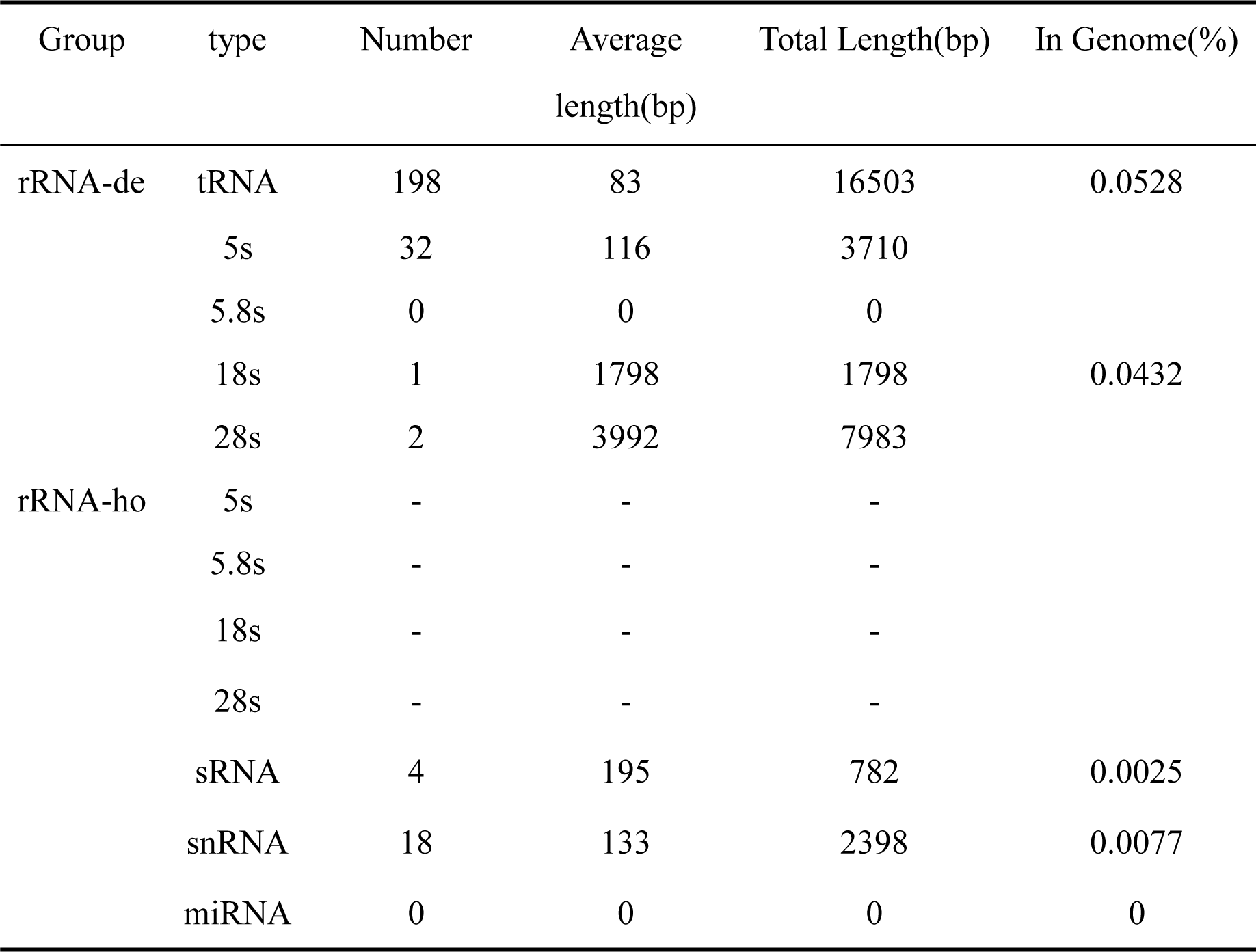
Statistical results of the *A. lentulus* non coding RNA after redundancy removal.

### 3.2. Genome annotation GO functional annotation

The statistical results of classification in GO entries for all genes of *A. lentulus* PWCAL1 is shown in figure 3. These genes are mainly enriched in metabolic process, catavytic activity, cell and cell part, which shows that *A. lentulus* has vigorous primary and secondary metabolic processes, as well as enzymatic catalytic activity to the external environment or itself. In addition, *A. lentulus* PWCAL1 is also posses abundant oxidation reduction, ATP binding and integral to membrane. GO entry analysis of gene with high impact SNPs and indels by Dos Santos et al showed that *A. lentulus* had enriched “nucleoside metabolic processes” and “glycosyl compound metabolic processes” [Dos Santos RAC et al.2020].

**Figure 3.**
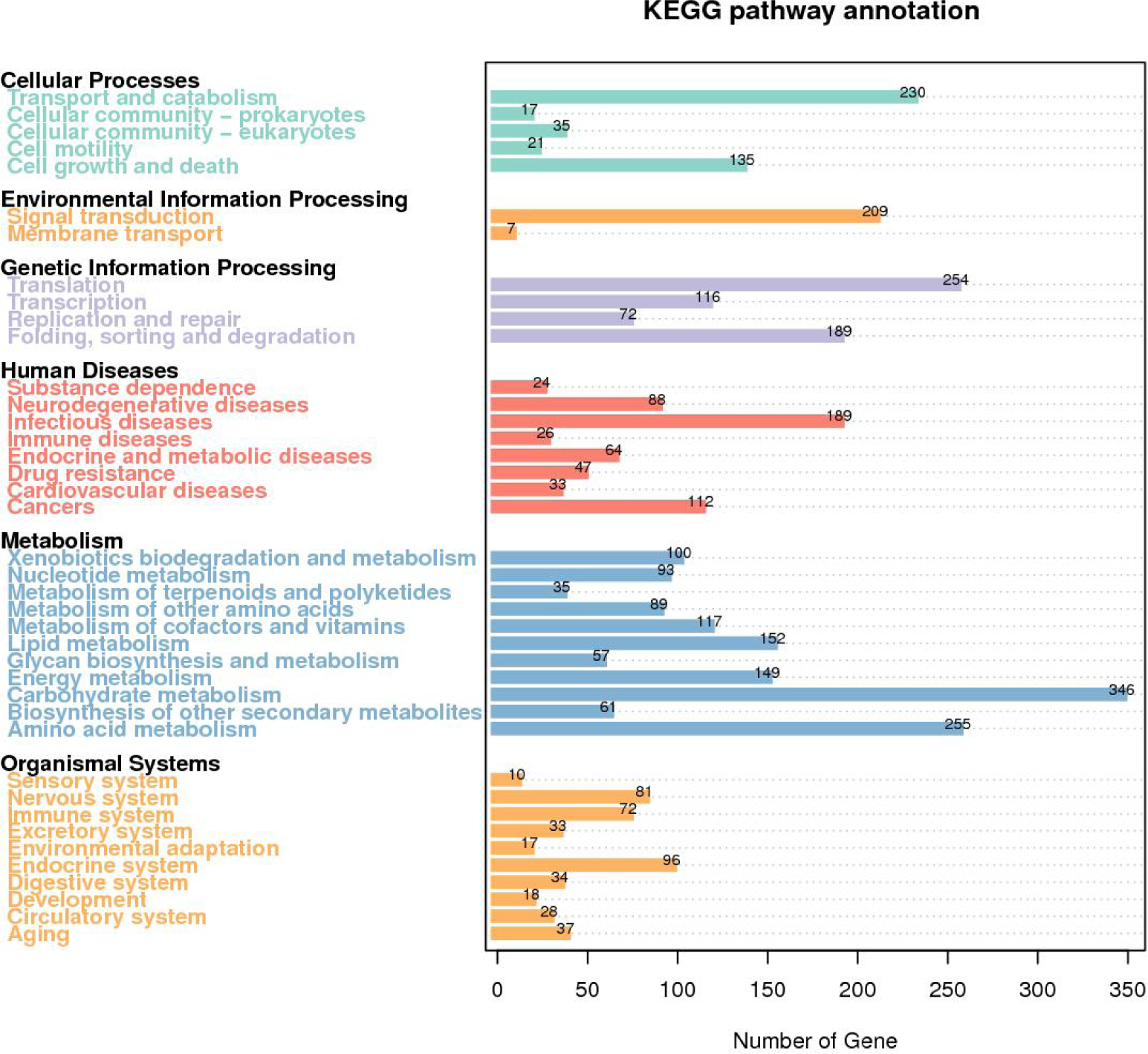
*A. lentulus* Gene Function Annotation of GO Functional Classification Map

#### KEGG functional annotation

The metabolic pathway classification of *A. lentulus* PWCAL1 in KEGG is shown in Figure 4. Among them, metabolic pathways are mainly enriched in Metabolism, which includes 11 annotated pathways , and the most 346 genes were annotated to Carbohydrate metabolism, which is the most basic metabolic activity of organisms and requires the formation of efficient Carbohydrate metabolism pathways synergistically by a large number of genes. Moreover, eight pathways were annotated in Human diseases, with the highest proportion of genes annotated to the infectious diseases pathway reaching 189, which fully demonstrates that *A. lentulus* is a pathogen causing invasive fungal diseases.

**Figure 4.**
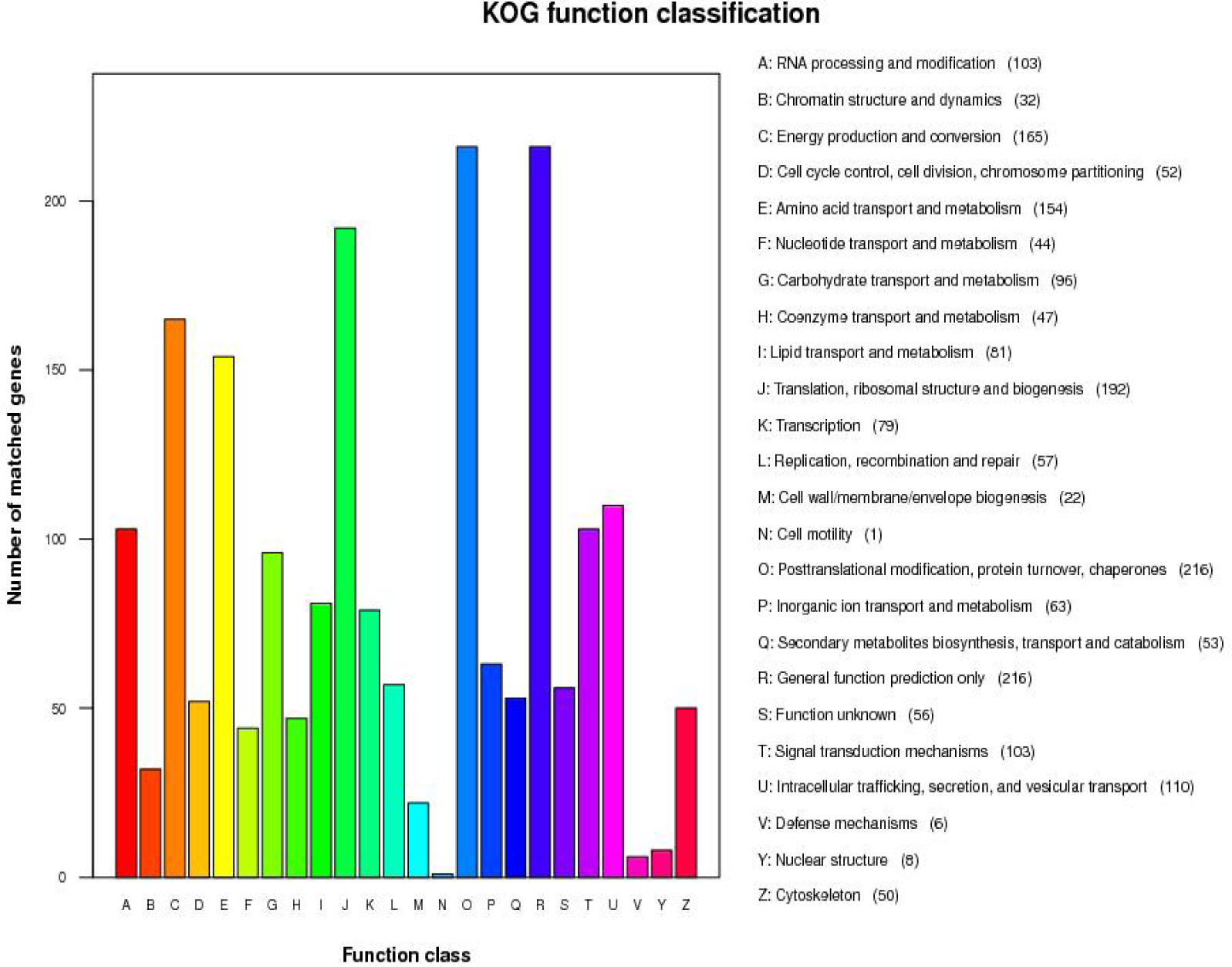
*A.lentulus* Gene Function Annotation of KEGG Metabolic Pathway Classification Map

The *A. lentulus* PWCAL1 genome was compared with the known metabolic pathway networks in the KEGG database to analyze the genes contained in different metabolic pathways and the distribution of genes involved in metabolic pathways, and 359 important KEGG signaling pathway maps were given, of which there were three main pathways related to energy metabolism (shown in Table 6). In the Autophagy-yeast pathway(map04138), 49 genes of *A. lentulus* may be involved in this metabolic pathway; in the Mitophagy-yeast pathway(map04139), 26 genes are involved in this pathway. Autophagy plays an important role in the pathogenicity of fungi. In the ABC transporters signaling pathway, there were 5 genes that are invloved which includes (A1024/K15628/−), (A1589/K15628/−), (A1885/K05663/−), (A3002/K05658/3.6.3.44) and (A0675/K05658/3.6.3.44). Among them, the A3002 gene and A0675 gene of *A. lentulus* PWCAL1 belong to ATP-binding cassette subfamily B (MDR/TAP), member 1, and A1024 and A1589 genes belong to ATP-binding cassette, subfamily D (ALD), peroxisomal long-chain fatty acid import protein, and both subfamilies are involved in antifungal resistanc. Studies have demonstrated that the extensive excretion of xenobiotics by ABC transporters is one of the main mechanism to develop clinical multi-resistance of pathogenic fungi, which also include Candida species[Moazeni M et al.2020;Jensen RH.2016;Tobin MB et al.1997;Pinjon E et al.2005].

**Table 6.**
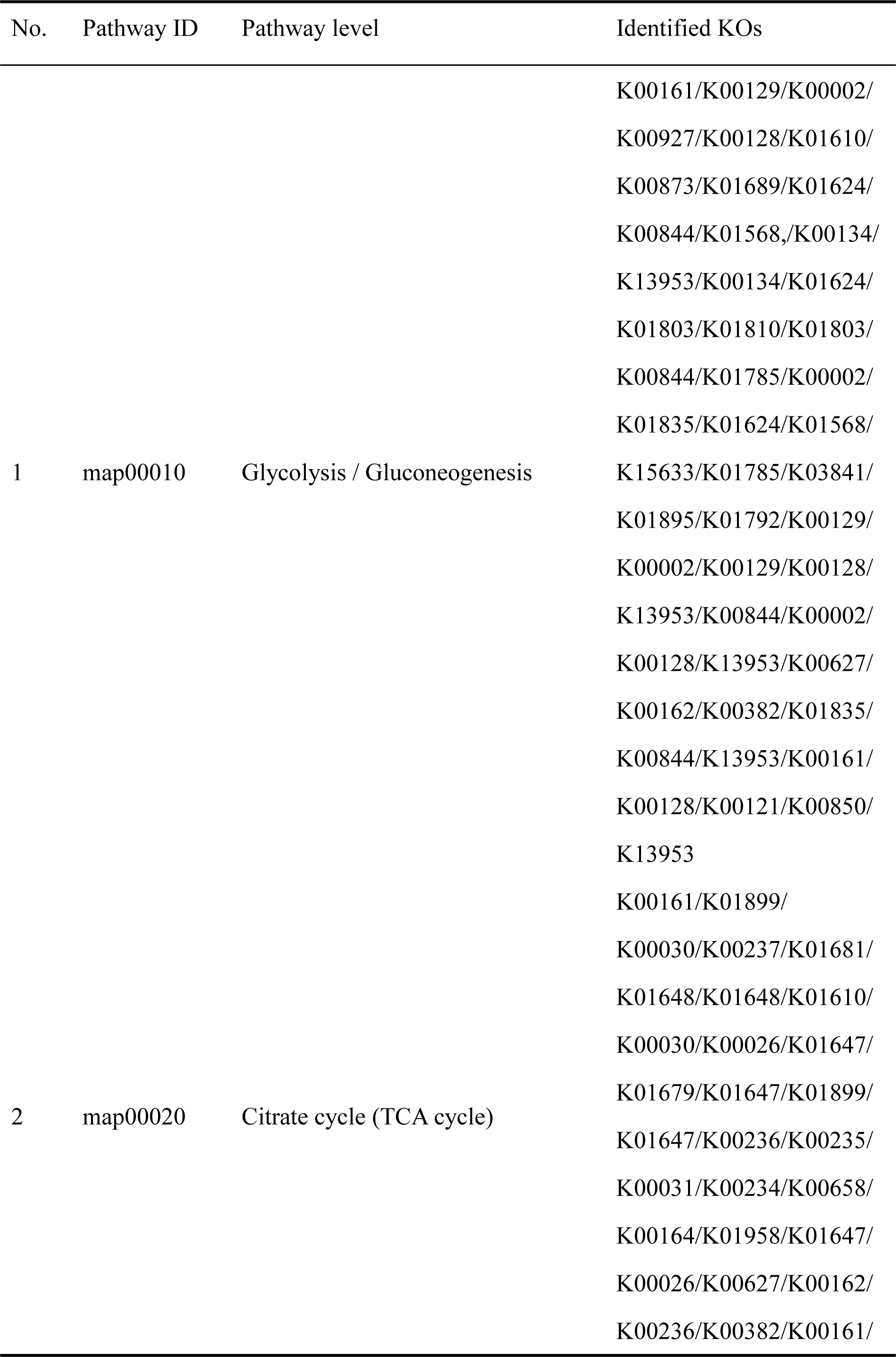
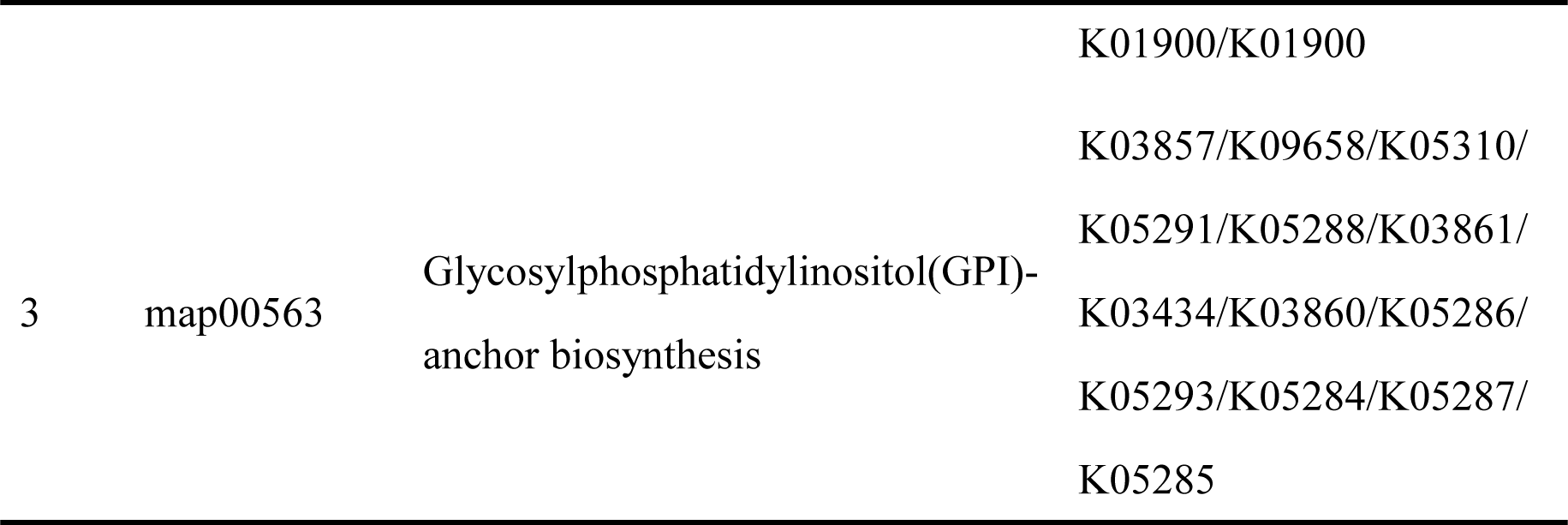
*A. lentulus* main pathway of energy metabolism.

#### KOG functional annotation

According to the statistic results of KOG functional annotation (see Figure 5), these genes are mainly enriched in post modification, protein turnover, chaperones (216 genes), post translational modification is a chemical modification that plays a key role in the functional proteome because they regulate activity and localize to interact with other cellular molecules such as proteins, nucleic acids, and lipids, and identification and understanding of these translational modification genes is essential in the study of cell biology and pathogenesis treatment and prevention. However, the less enriched genes were Nuclear structure (8 genes), Defense mechanisms (6 genes), and cell motility (1 gene). In addition, there were 56 unknown functional genes, which may be unique functional genes of *A. lentulus* PWCAL1. From the above results, it can be shown that in microbial genomes, different types of genes are quantitatively focused, and it is related to the complexity of executive function and the embodiment of the adaptability formed by organisms in long-term evolution.

**Figure 5.**
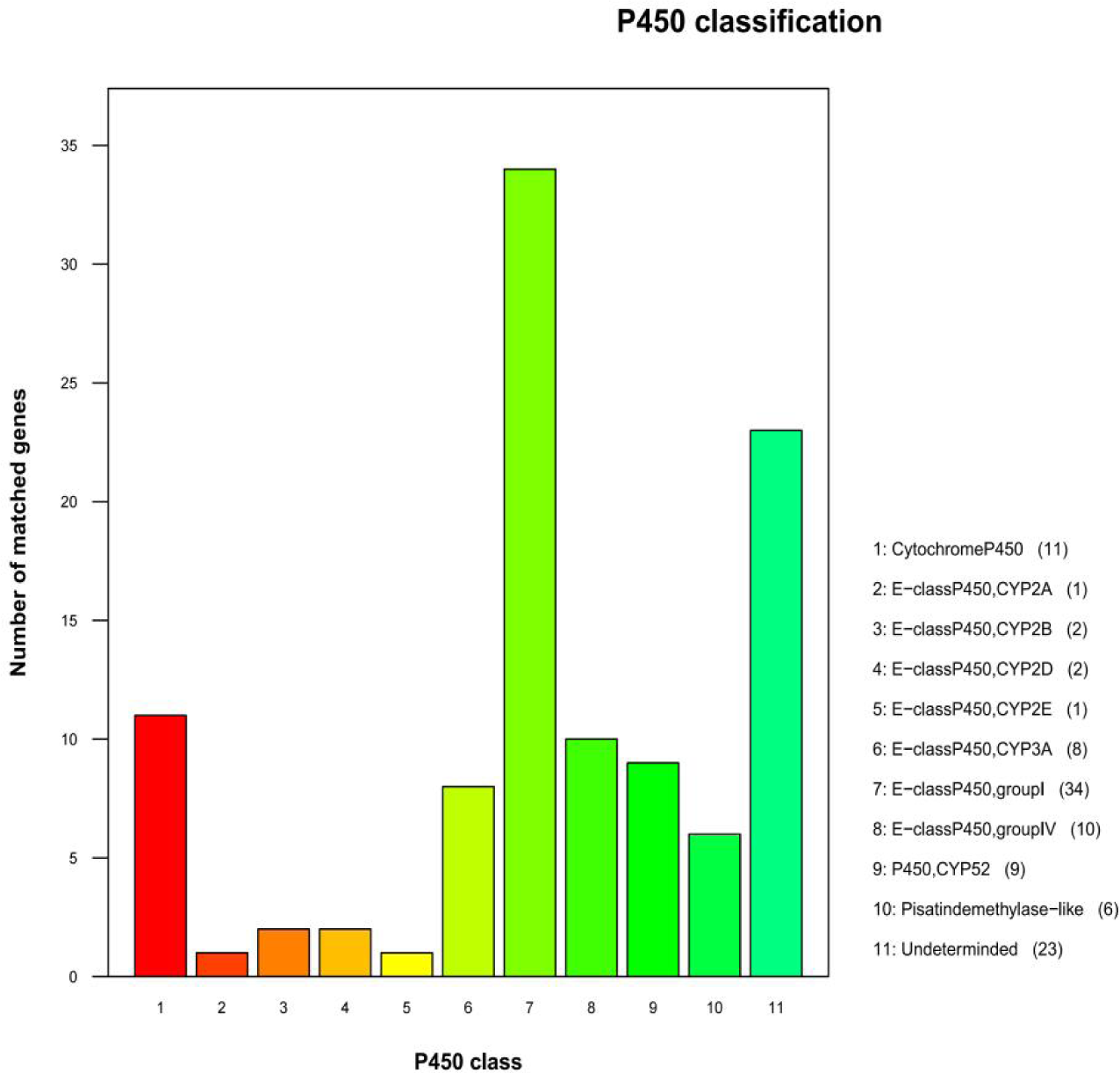
*A.lentulus* Gene Function Annotation of KOG Functional Classification Map

#### TCDB annotation

Based on the Transporter Classification Database (TCDB) annotation results, the first level functional classification of *A. lentulus* PWCAL1 transporters is mainly enriched in Electrochemical Potential-driven Transporters (157 genes). Transporter class II functional classification was enriched mainly in Porters (uniporters, symporters, antiporters), with a total of 156 genes.

#### Carbohydrate-Active enZYmes Database (CAZy) Database annotation Secreted

CAZymes are essential for the biological activity of fungi. Based on the CAZy database, a total of 491 CAZymes were identified, which includes 272 Glycoside Hydrolases, 87 Glycosyl Transferases, 57 Carbohydrate-Binding Modules, 39 Auxiliary Activities, 21 Carbohydrate Esterases, and 15 Polysaccharide Lyases. Carbohydrate binding domain is a non-catalytic domain that folds into a specific three-dimensional spatial structure and has the function of binding carbohydrates. Carbohydrate binding domain can improve the activity of carbohydrate active enzyme by binding to substrate.

#### P450 Database annotation

As shown in Figure 6, functional classification showed that it belongs to 11 P450 gene families, mainly concentrated in the group I and IV family of E-classP450, of which the A5653 gene of *A. lentulus* PWCAL1 was annotated as the CYP51 gene and involved in the mechanism of azole resistance in fungi.

**Figure 6.**
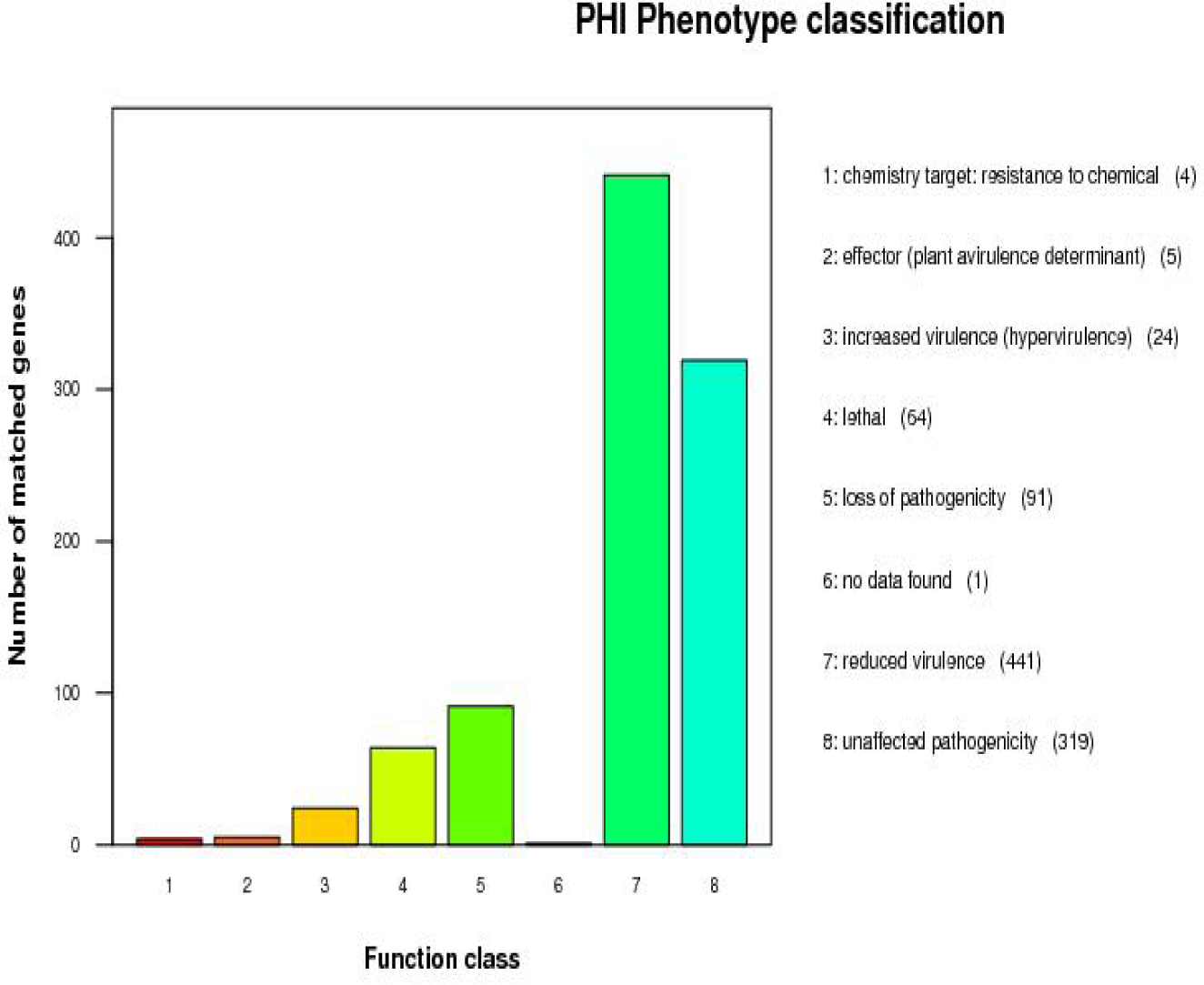
*A. lentulus* P450 Database Annotation Classification

#### Pathogen Host Interactions (PHI) Database annotation

The statistics of the number of PHI phenotype mutant genes in *A. lentulus* PWCAL1 pathogens are shown in Figure 7, and the mutation types of this species are mainly concentrated in reduced virulence and unaffected virulence, which are consistent with the aligned strains *A. lentulus* (IFM54703, CNM-CM8927), *A. fumigatiaffinis* (CNM-CM5878), and *A. fumigatus* (Af293, A1163). Deletion of these genes has been shown to result in decreased virulence of the species and loss of pathogenicity in mutant strains PHI annotation results showed that the top three virulence factors were *Fusarium graminearum* (259 genes), *Magnaporthe oryzae* (150 genes) and *A. fumigatus* (134 genes). The diseases caused were mainly Fusarium ear blight (212 genes), Rice blast (149 genes) and Invasive aspergillosis (106 genes). It can be seen that these virulence factors not only cause plant diseases, but also cause human infections. These virulence factors are abundantly expressed in *A. fumigatus* and play a role in the pathogenesis of Invasive pulmonary aspergillosis. Transcription factor may perform transcriptional activation and regulation in the mechanism of *Aspergillus* resistance.

**Figure 7.**
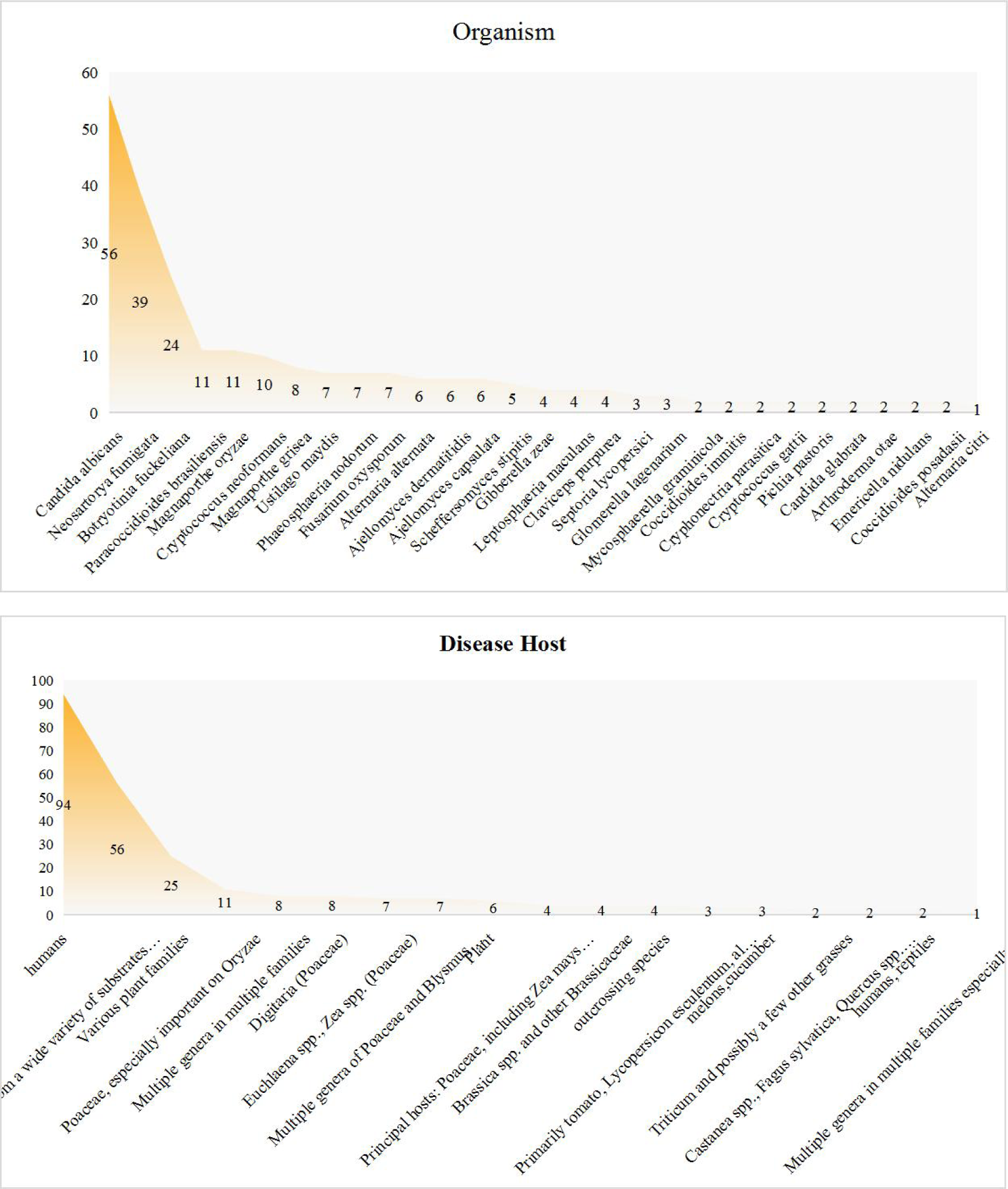
*A.lentulus* Gene Function Annotation Pathogen of PHI Phenotypic Mutation Type Distribution Map

Notably, in PHI, we identified four virulence genes (A1704, A2802, A2958, A3108) belongs to abcB and one virulence gene (A4203) belongs to abcA in *A. lentulus* PWCAL1, all of which are ABC transporter-genes which can cause Invasive pulmonary resistance-aspergillosis, and its mutant phenotypes are annotated to Multidrug resistance cdr1 (A1704, A2802, A3108, A4203) and cdr2 (A2958) in the TCDB database, respectively. The abcA and abcB proteins of *A. fumigatus* have been shown to share high similarity with the pdr5 sequence encoded by the ABC transporter of *S. cerevisiae*, with both proteins localized in the plasma membrane and overproduced in azole resistant yeasts[Egner R et al.1995;Paul S et al.2013]. Paul et al have studied the resistance and virulence of abcA and abcB proteins in *A. fumigatus* and results have demonstrated that abcB (cdr1B) as an important participant in the biology of drug resistance in *A. fumigatus*, Virulence assays implicated abcB as a possible factor required for normal pathogenesis of the fungus, and overproduction of abcA also yielded increased azole resistance[Paul S et al.2013]. It also suggests that, like *A. fumigatus*, the above virulence genes present in *A. lentulus* PWCAL1 may have the same gene function.

#### Database annotation of fungal virulence factors (DFVF)

Alignment of the *A. lentulus* PWCAL1 genome with virulence factors from the database of fungal virulence factors identified a total of 259 interacting factors, and the result shown that there were 167 gene annotated to the corresponding functions. Among them, the most annotated organism was *Candida abbicans* and the most common host types are humans. The type of disease host keys are vertebrata animal and herb. The mainly caused disease-key are invasive candidal disease(56,22%) (see Figure 8A-C), which was consistent with the annotation results of the aligned strains *A. lentulus* (IFM54703, CNM-CM8927), *A. fumigatifumis* (CNM-CM5878), and *A. fumigatus* (Af293, A1163). These results indicate that the pathogenicity of different species in the *A. fumigatus* complex has common characteristics.

**Figure 8.**
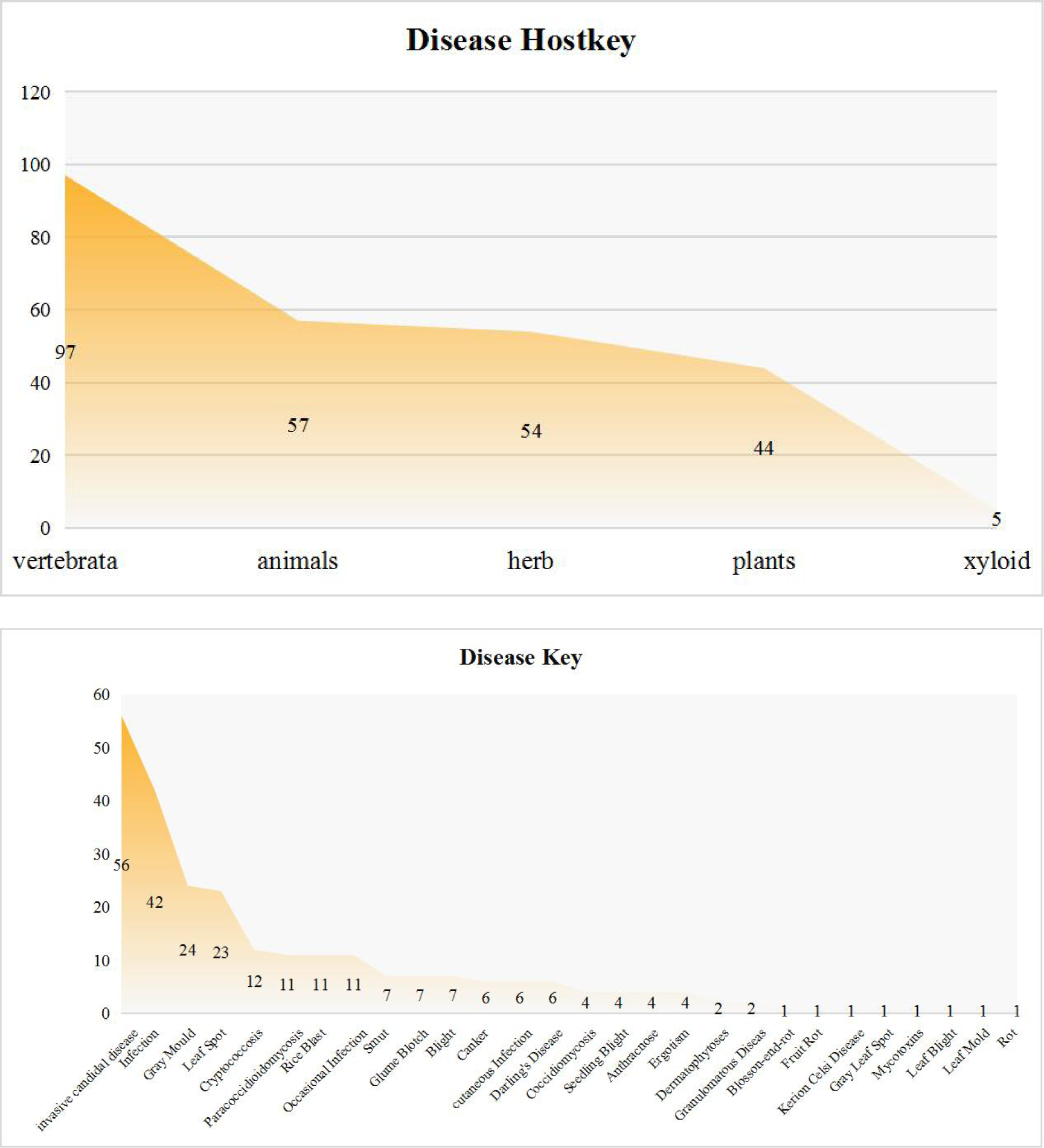
(A-C). Comparison results of virulence factors between *A. lentulus* genome and database of functional virus factors

### 3.3. Comparative genomic analysis (Core Gene and Specific Gene)

The complete genome sequences of four closely related strains, *A. lentulus* (IFM54703), *A. lentulus* (CNM-CM6936), *A. fumigatiaffinis* (CNM-CM6457), and *A. fumigatus* (Af293), were downloaded from NCBI Genome and analyzed for core genes and specific genes together with the complete genome sequences of *A. lentulus* PWCAL1. According to Wenckebach plot analysis of Pan Gene homology relationship (see Figure 9), the number of all non-redundant Pan Genes between *A. lentulus* PWCAL1 and the four model strains was 11,102, including 5456 Core Genes. Specific Gene numbers of each strain were 806 *A. fumigatus* (Af293), 610 *A. fumigatiaffinis* (CNM-CM6457), 171 *A. lentulus* PWCAL1, 129 *A. lentulus* (CNM-CM6936), and 96 *A. lentulus* (IFM54703). Among the three *A. lentulus* strains, *A. lentulus* PWCAL1 had the highest number of Specific Gene.

**Figure 9.**
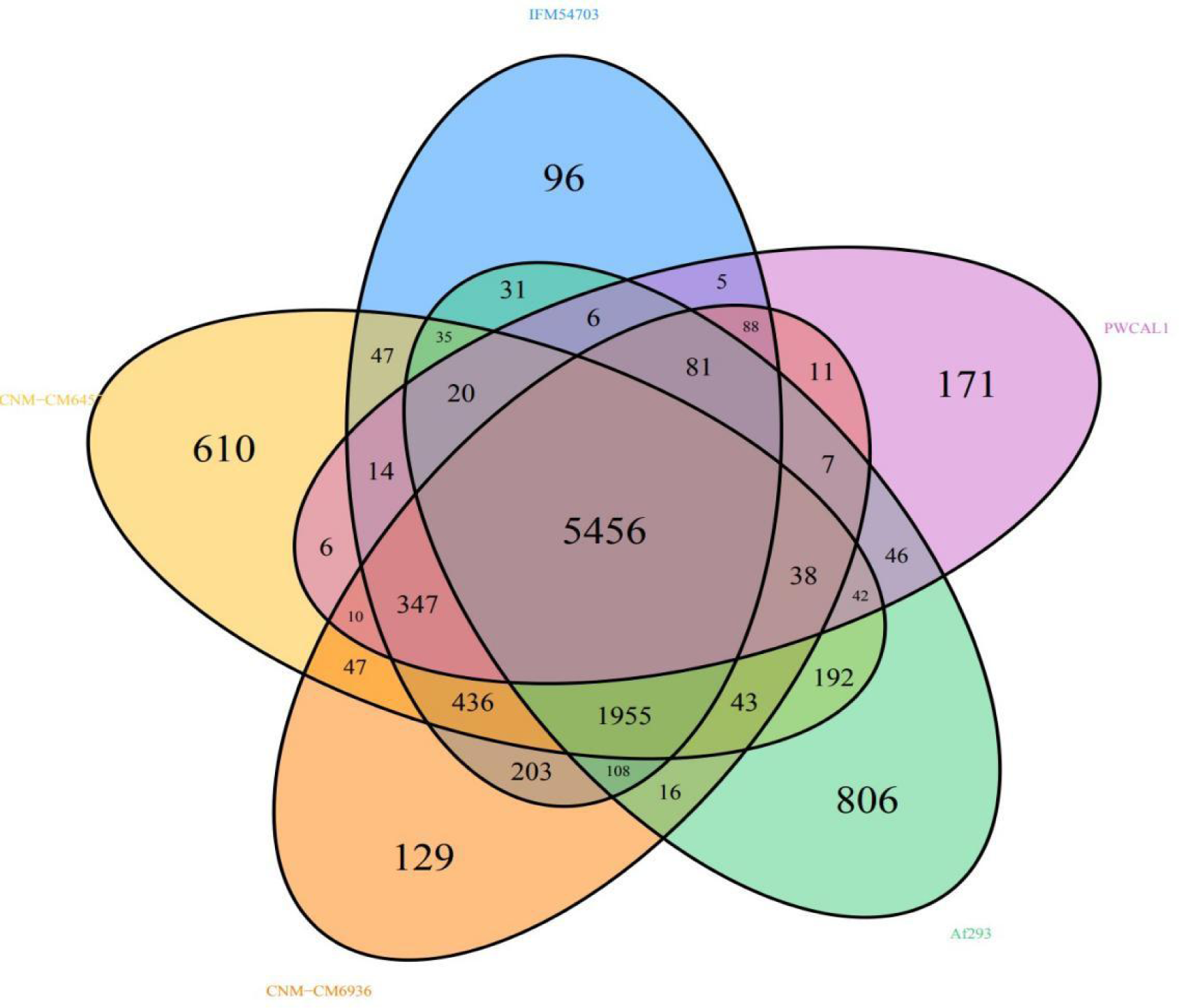
Wen’s diagram of the Pan Gene homologous relationship between core genes and specific genes

#### Core genes and Specific genes in KEGG and KOG Database annotation

The 5,456 core genes of five aligned *A. lentulus* strains were annotated to the KEGG database, and it was observed that these genes were mainly enriched in Human Diseases, Metabolism, Organismal Systems, Cellular Processes, Gene (see Figure 10A). In the KOG database annotation, these genes were mainly enriched in general function prediction only, posttranslational modification, protein turnover, chaperones, Translation, ribosomal structure and biogenesis (see Figure 10B).

**Figure 10A.**
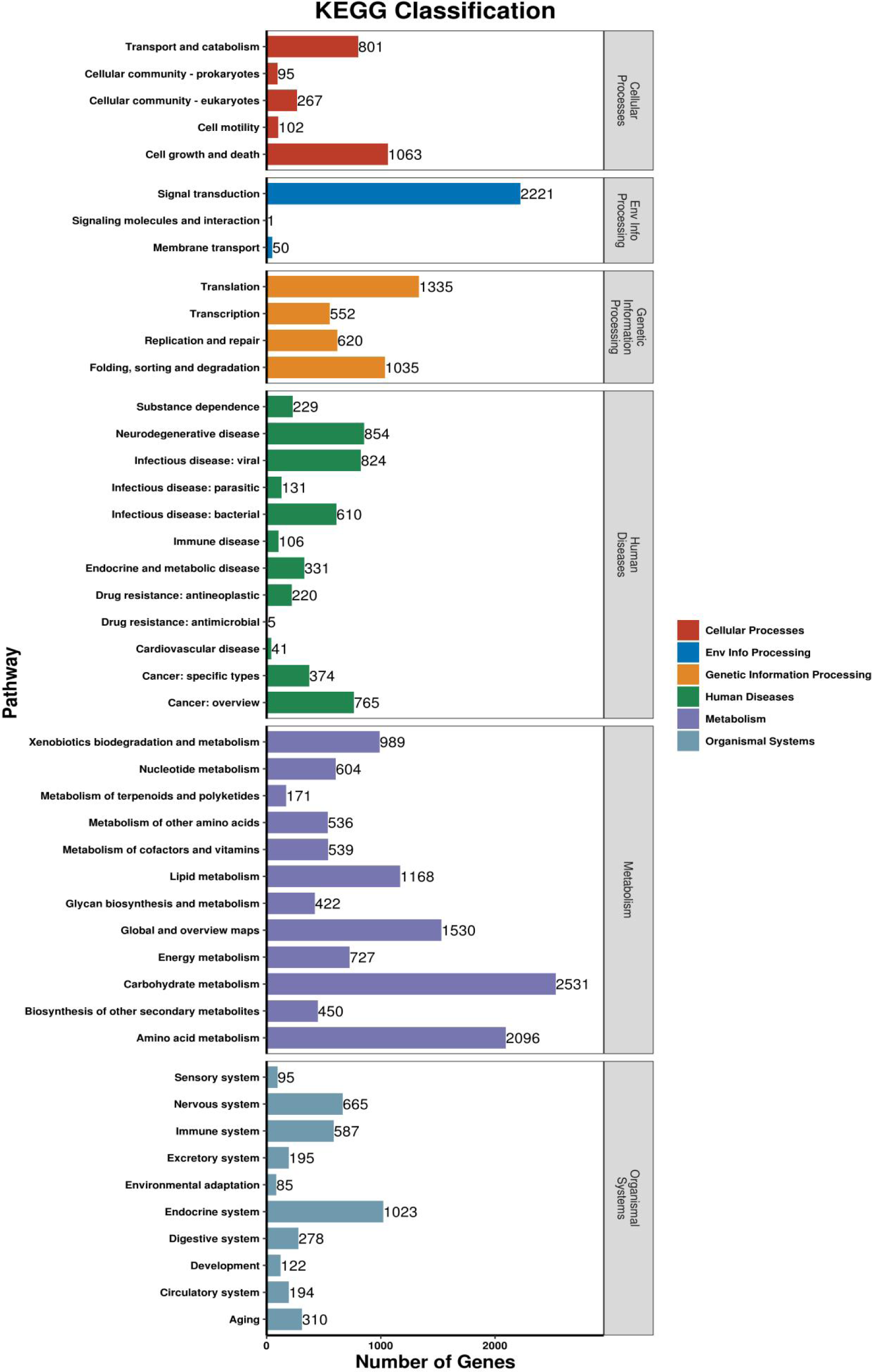
Classification of KEGG Functional Annotations of 5456 Core Genes

**Figure 10B.**
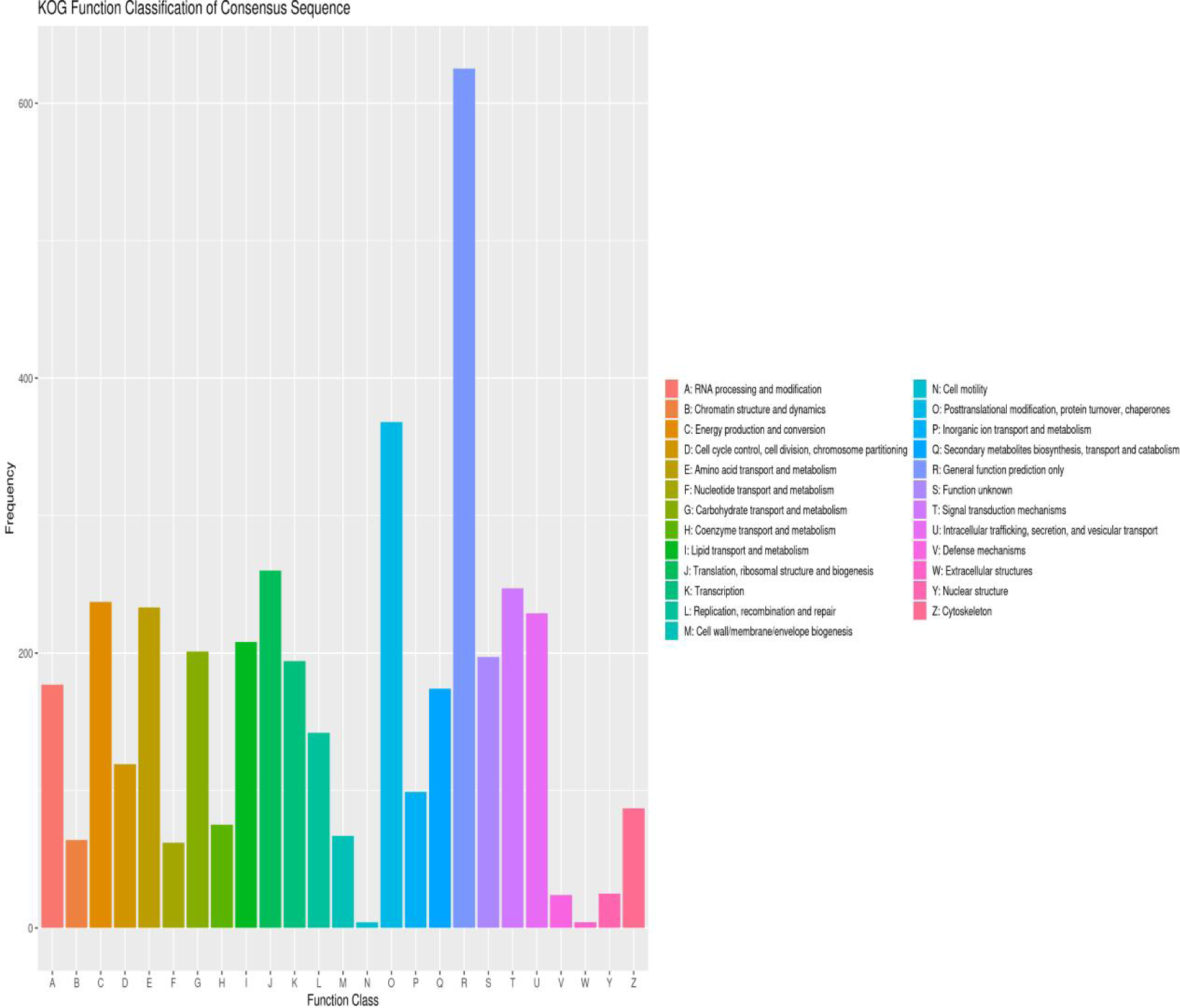
KOG Functional Annotation Classification of 5456 Core Genes

The 171 specific genes of A. lentulus PWCAL1 were annotated to the KEGG database (see Figure 10C) and these genes were mainly enriched in Carbohydrate metabolism, Cell growth and death. The 129 specific genes of *A. lentulus* (CNM-CM6936) were mainly enriched in Carbohydrate metabolism, Amino acid metabolism, Signal transduction. The 96 specific genes of *A. lentulus* (IFM54703) were mainly enriched in Amino acid metabolism.

**Figure 10C.**
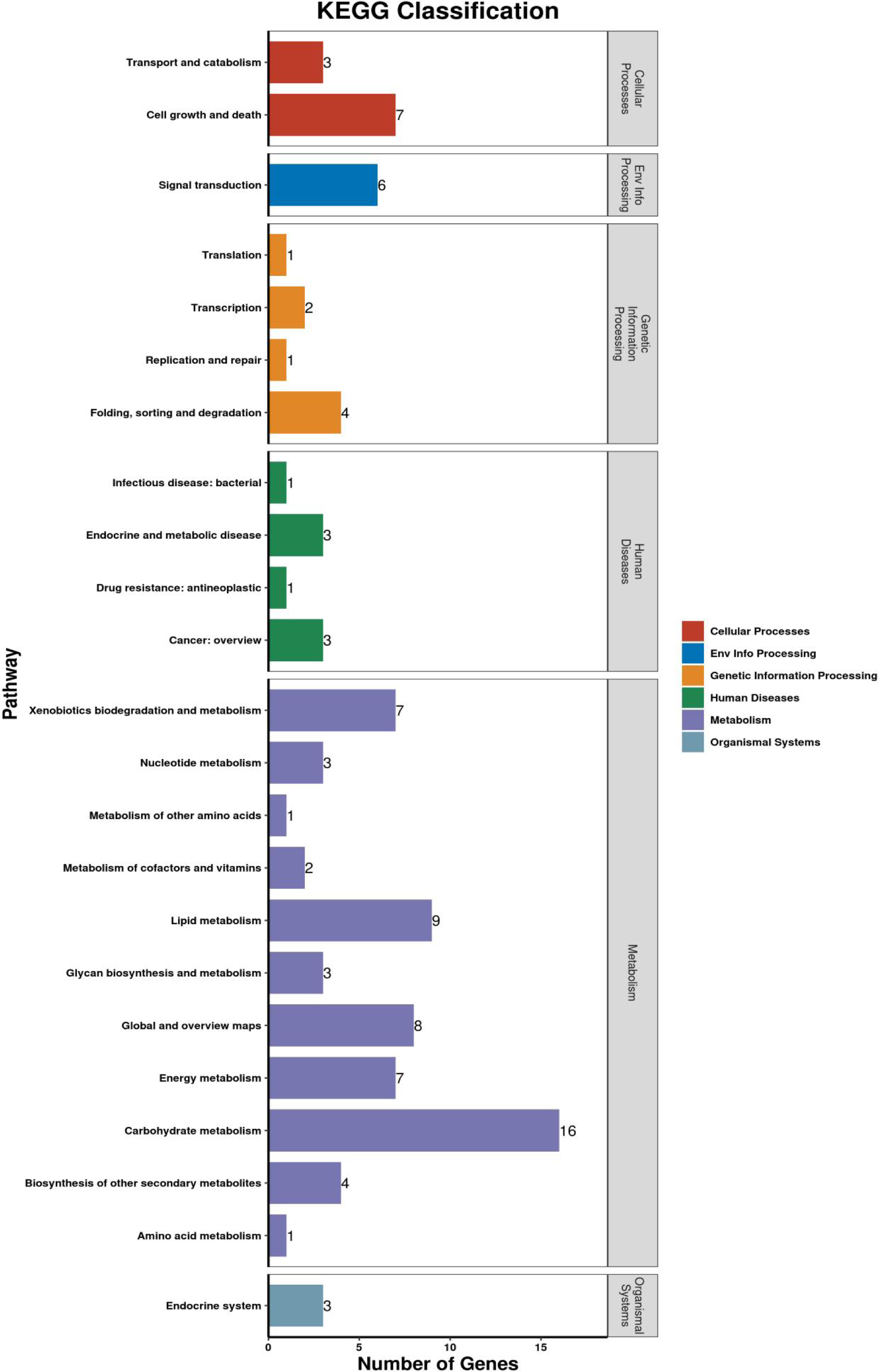
Classification of the *A. lentulus* 171 specific gene in KEGG functional annotations

The 171 specific genes of *A. lentulus* PWCAL1 were annotated to the KOG database (see Figure 10D). These genes are mainly enriched in General function prediction only, Signal transduction mechanisms. The 129 specific genes of *A. lentulus* (CNM-CM6936) were mainly enriched in Energy production and conversion, Secondary metabolites biosynthesis, transport and catabolism. The 96 specific genes of *A. lentulus* (IFM54703) were mainly enriched in General function prediction only.

**Figure 10D.**
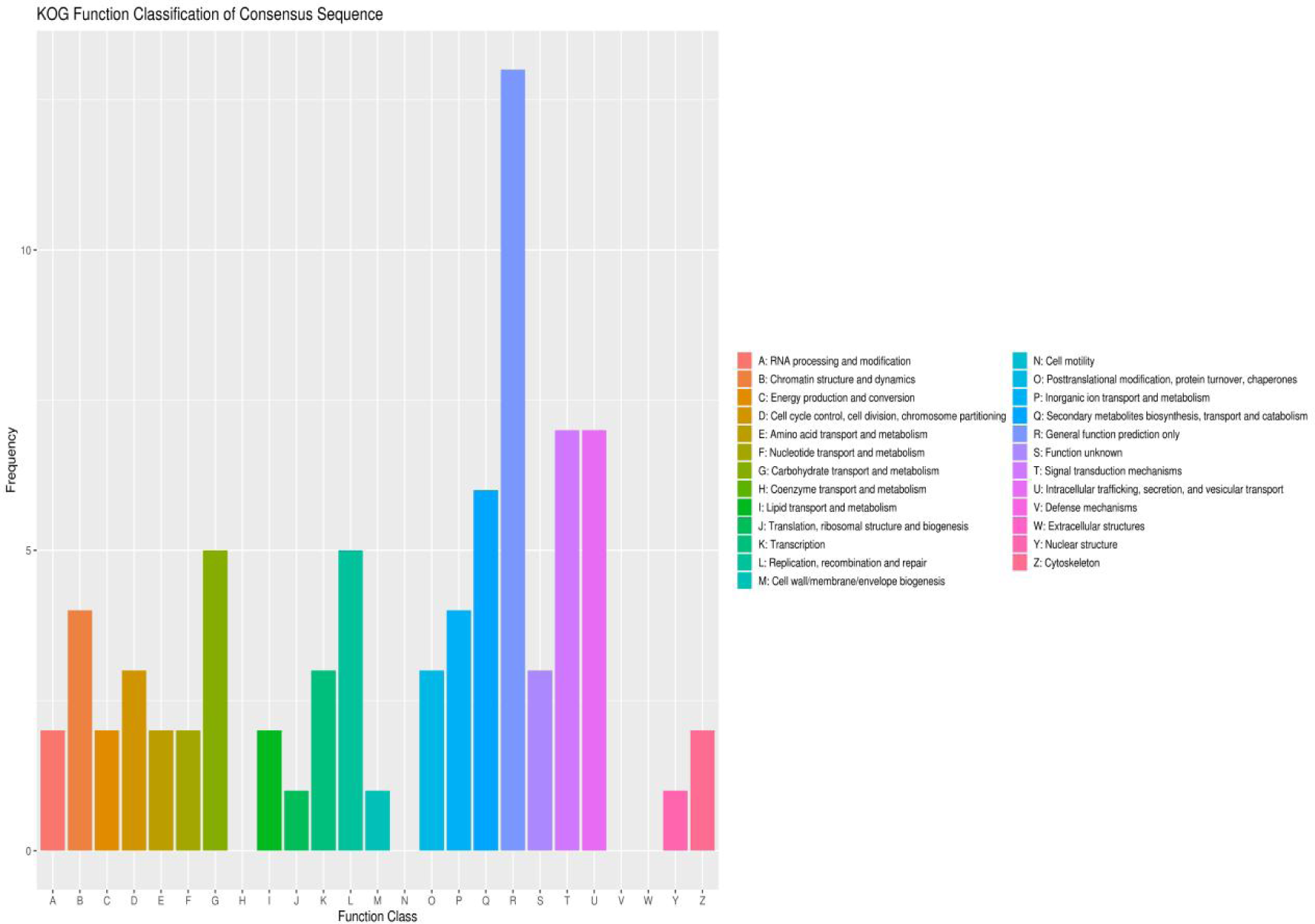
Classification of the *A. lentulus* 171 specific gene in KOG functional annotations

### 3.4. Screening analysis of *A.lentulus* PWCAL1 resistance genes using comparative genomics study

#### (1) Mutation analysis of CYP51A gene region

**① Screening analysis of point mutation**

We performed blastp alignment of *A. lentulus* PWCAL1 genomic data with Mycology Antifungal Resistance Database [Nash A et al.2018]. Alignment results showed that the A5653 gene of this strain was aligned to the Cyp51A gene of Af293 in the database – Afu4g06890 (with the F46Y/M172V/N248T/D255E/E427K mutation combination). The mutations present in the A5653 gene on the alignment intersected with the F46Y and N248T mutations recorded in the database, corresponding to resistance information of Itraconazole and Posaconazole, from which we speculated that *A. lentulus* PWCAL1 had the F46Y/N248T mutation combination on the Cyp51A gene. Since 2001, a combination of Cyp51A amino acid substitutions (F46Y, M172V, N248T, D255E, and E427K) has been increasingly reported worldwide [van der Linden JW et al.2015; Pérez-Cantero A et al.2020]. These mutations emerged in azole susceptible or resistant A. fumigatus strains either individually or in combination and belong to specific missense Cyp51A mutations. Garcia-Rubio showed that A. fumigatus carrying the CYP51A-F46Y mutation had a high MIC value for azoles due to obstruction of the substrate channel for drug action [Garcia-Rubio R et al.2018].

However, some investigators believe that combinations of these mutations are not associated with azole resistance [Mellado E et al.2007], as MIC values for azoles in strains with combinations of these mutations generally remain below acceptable thresholds [Verweij PE et al.2009].It is therefore controversial whether these different combinations of mutations enforce azole resistance in *Aspergillus*.

Studies on the mechanisms of *A. lentulus* resistance are very limited. Mellado et al reported that introduction of *A. lentulus* CYP51A(Alcyp51A) into *A. fumigatus* resulted in azole resistance and that the strain partially lacking Alcyp51A became susceptible to azole drugs[Mellado E et al.2011]. Tateno et al successfully introduced the entire *A. fumigatus* cyp51A gene into the cyp51A locus in *A. lentulus* using the CRISPR/Cas9 genome-editing system. The *A. lentulus* strains harboring *A. fumigatus* cyp51A showed reduced MIC for itraconazole and voriconazole compared with the parent strain[Tateno M et al.2022]. However, none of these studies clarified which specific mutation sites were present on the *A. lentulus* CYP51A gene. However, the F46Y/N248T mutation combination on *A. lentulus* CYP51A gene was detected for the first time by whole gene sequencing in this study. We confirmed that the resistance of *A. lentulus* to azoles is partly responsible for the CYP51A gene by whole genome sequencing results and consistent with previously reported CYP51A gene replacement studies.

**② Screening analysis of TR (tandem repeat) -mediated mutations**

CYP51A of *A. fumigatus* strain F13560 was used as the reference sequence (GenBank: JX283443.1) to blastn alignment with the genome sequence of the under tested *A. lentulus* PWCAL1 strain, and this gene sequence of *A. lentulus* PWCAL1Contig6 (2354792-2356409) was aligned, that is, the gene sequence of this region was the CYP51A gene of interest on the under tested strain. However, comparing the gene sequence of this region with the mutation information recorded in the MARDy database, there were no intersection sites, indicating that no amino acid mutations were detected in *A. lentulus* PWCAL. Meanwhile, no tandem repeats were detected in the gene sequences of this region when repeat prediction analysis was performed with TRF (Tandem Repeats Finder, Version 4.07b) software. This indicate that there is no TR associated mutation in the *A. lentulus* PWCAL1 CYP51A gene. Chong et al showed that the AsperGennius resistance PCR did not detect the TR_34_ target in *A. lentulus*(n=7) and *A.felis*(n=5) in contrast to *A. fumigatus* [Chong GM et al.2016]. The results of the present study have demonstrated that whole gene sequencing alignment were consistent with Chong ’s PCR amplification study, and we speculated that *A. lentulus* may not have a TR-related mutation resistance mechanism, and this phenomenon is whether *A. lentulus* generally has or is unique to individual strains requiring further expansion of specimen size for validation.

Therefore, based on the detection of the CYP51A-F46Y/N248T mutation combination in the *A. lentulus* PWCAL1 strain, we suggest that resistance to azoles in *A. lentulus* may be mediated in part by mutations in the CYP51A gene.

#### (2) ABC transporters subfamily prediction

In the TCDB annotated secondary functional classification, 19 genes of *A. lentulus* PWCAL1 were annotated to The ATP-binding Cassette (ABC) Superfamily (see Table 7), of which 12 genes were annotated to fungal multidrug-resistant ABC transporters, among which, A1589 belongs to the ALDp subfamily, A2265/AtrB, A6788/AtrB, A0601/AtrB, A1704/cdr1, A2802/cdr1, A3108/cdr1, A4482/cdr1, A4203/cdr1, A2958/cdr2 belongs to the PDR subfamily, and A3002/mdr1, A0675/mdr1 belongs to the MDR subfamily. This may suggest a very different capability for drug efflux, and we speculated that differences in the uptake of azoles or differences in intracellular azole concentrations within *A. lentulus* due to increasced expression of efflux pumps such as ABC transporters may play a role. In recent study, Moazeni et al have found that overexpression of efflux pump genes is an alternative mechanism in voriconazole resistant *A. fumigatus* isolates without relative mutations in CYP51A[Moazeni M et al.2020].

**Table 7.** Identification of ABC transporters and their subfamily classification in 9 strains of *Aspergillus fumigatus* complex.

| Species | numbers of ABC | PDR |  |  |  | MDR |  |  |  | ALDp |
| --- | --- | --- | --- | --- | --- | --- | --- | --- | --- | --- |
|  |  | cdr1 | cdr2 | AtrB | AtrF | mdr | mdr1 | mdr2 | mdr3 |  |
| *A.lentulus-PWCAL1* | 12/19 | A1704<br>A2802<br>A3108<br>A4203<br>A4482 | A2958 | A2265<br>A6788<br>A0601 | / | / | A3002<br>A0675 | / | / | A1589 |
| *A.lentulus-IFM54703* | 13/34 | / | / | / | GAQ03337.1<br>GAQ03595.1<br>GAQ12074.1 | / | GAQ03747.1<br>GAQ08512.1<br>GAQ09078.1<br>GAQ09976.1 | GAQ08155.1 | GAQ04138.1<br>GAQ06261.1<br>GAQ08677.1<br>GAQ09205.1<br>GAQ10522.1 | / |
| *A.lentulus-CM-8927* | 14/34 | / | / | KAF4207593 | KAF4201478.1<br>KAF4202847.1<br>KAF4205113.1 | / | KAF4206090.1<br>KAF4207019.1<br>KAF4207242.1<br>KAF4207827.1 | KAF4201033.1 | KAF4202325.1<br>KAF4204142.1<br>KAF4206915.1<br>KAF4207341.1<br>KAF4209521.1 | / |
| *A.lentulus-CM-6936* | 14/33 | / | / | KAF4170081.1 | KAF4159563.1<br>KAF4162838.1<br>KAF4164632.1 | / | KAF4167062.1<br>KAF4167350.1<br>KAF4167461.1<br>KAF4167584.1 | KAF4165326.1 | KAF4160620.1<br>KAF4163223.1<br>KAF4163585.1<br>KAF4165529.1<br>KAF4169109.1 | / |
| *A.udagawae-IFM46973* | 14/38 | / | / | XP_043143397 | XP_043145569.1<br>XP_043145840.1<br>XP_043149814.1 | / | XP_043145999.1<br>XP_043147149.1<br>XP_043147672.1<br>XP_043147757.1 | XP_043148163.1 | XP_043142908.1<br>XP_043144526.1<br>XP_043147576.1<br>XP_043148949.1<br>XP_043150390.1 | / |
|  |  | cdr1 | cdr2 | AtrB | AtrF | mdr | mdr1 | mdr2 | mdr3 |  |
| *A.fumigatus-Af293* | 15/32 | / | / | XP_746352.2<br>XP_748461.2 | XP_746942.1<br>XP_747642.1<br>XP_750693.1 | XP_746882.1<br>XP_747730.1 | XP_747768.1<br>XP_751419.1<br>XP_754025.1 | XP_751826.1 | XP_748261.1<br>XP_752803.1<br>XP_753111.1<br>XP_755847.1 | / |
| *A.fumigatiaffinis-CNM.CM8980* | 18/39 | / | / | KAF4246624.1<br>KAF4249070.1 | KAF4238890.1<br>KAF4241484.1<br>KAF4251827.1 | KAF4233685.1 | KAF4245202.1<br>KAF4245521.1<br>KAF4250115.1<br>KAF4250835.1 | KAF4246531.1 | KAF4237305.1<br>KAF4246959.1<br>KAF4247240.1<br>KAF4247244.1<br>KAF4248511.1<br>KAF4248680.1<br>KAF4250207.1 | / |
| *A.fumigatiaffinis-CNM-CM6457* | 17/40 | / | / | KAF4235340.1 | KAF4225881.1<br>KAF4229335.1<br>KAF4239617.1 | KAF4228894.1 | KAF4222052.1<br>KAF4224161.1<br>KAF4237646.1<br>KAF4241268.1 | KAF4225042.1 | KAF4222028.1<br>KAF4223214.1<br>KAF4224505.1<br>KAF4224509.1<br>KAF4231832.1<br>KAF4238247.1<br>KAF4238738.1 | / |
| *A.viridinutans-FRR.0576* | 10/25 | / | / | A5724<br>A6341 | A3536<br>A4064 | / | A4172<br>A5044<br>A6519 | A5321 | A4674<br>A1495 | / |

To further understand the ABC transporters implicated in resistance and their taxonomic characteristics in other *A. fumigatus* complex species, we predicted their ABC transporter subfamilies by TCDB database annotation (see Table 7 and Figure 11). A total of 9 strains (including 4 strains of *A. lentulus*, 1 strain of *A. udagawae1*, 1 strain of *A. fumigatus*, 2 strains of *A. fumigatiaffinis*, and 1 strain of *A. viridinutans*) were selected to construct a phylogenetic tree of ABC protein subfamilies (see Figure 11), and these selected proteins had two characteristic topologies: Transmem binding brane domain (TMD) and Nucleotide binding domain (NBD). Among them, *A. fumigaffinatiis*-CNM.CM8980 possesses the most ABC protein members (18), while *A.Viridinutans*-FRR.0576 possessed the fewest ABC proteins (10). Based on the phylogenetic tree analysis, ABC proteins annotated in 9 strains were divided into three main subfamilies: (I) pleiotropic drug resistance (PDR), (II) multidrug resistance (MDR), and (III) adrenoleukodystrophy protein (ALDp). Non-predicted proteins were classified as unknown. Of these three subfamilies, none of the complex species were annotated to ALDp except for one protein of *A. lentulus* PWCAL1 annotated to ALDp. In the MDR subfamily, mdr3 was the most annotated, followed by mdr1, and finally mdr2 in these nine strains, but mdr2 and mdr3 were not observed in *A. lentulus* PWCAL1, which was significantly different from the other eight *A. fumigatus* complex species. In the PDR subfamily, AtrF was the most annotated in nine strains, followed by AtrB, but *A. lentulus* PWCAL1 was not annotated to AtrF. In CDR, only *A. lentulus* PWCAL1 was annotated to cdr1 and cdr2, while other complex species were not annotated to cdr1 and cdr2, and some previous studies have reported that cdr1 of ABC protein contributes more to azole resistance than cdr2 in clinically isolated azole-resistant C. albicans. The above results suggest that the distribution of different subfamilies of ABC transporters predicted is different among different *A. fumigatus* complex species.

**Figure 11.**
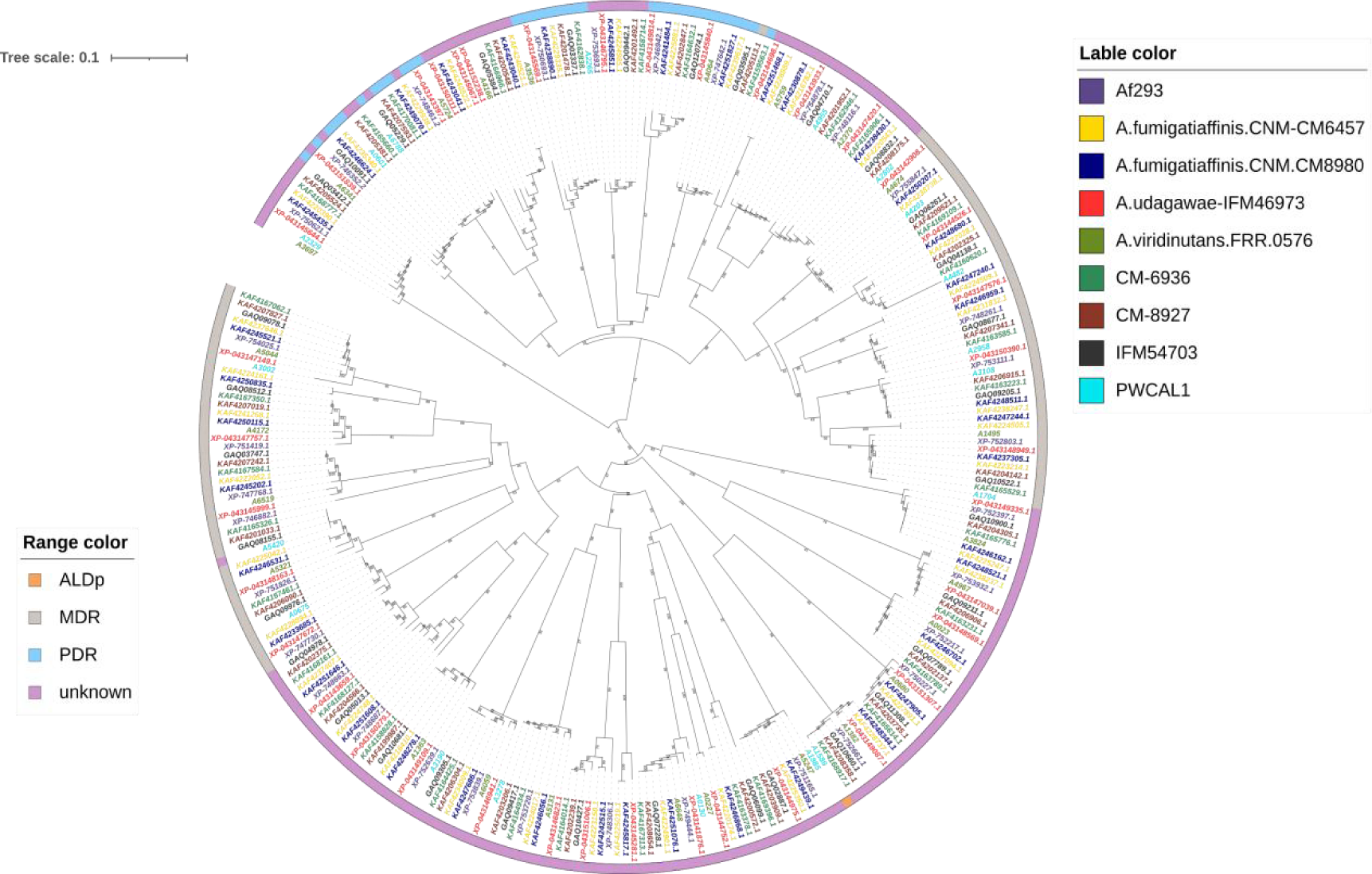
Comparison of ABC proteins in different *Aspergillus fumigatus* complex species. Note:Phylogenetic tree of the ABC proteins of nine *Aspergillus fumigatus* complex species.The numbers on the branches indicate the percentage of bootstrap support from 1000 replicates. The ABC subfamilies are identified based on known subfamilies in fungal species.

Among the four *A. lentulus* strains, MDR subfamily classification showed that except *A. lentulus* PWCAL1, mdr3 and mdr1 were mainly annotated in the other three strains, while mdr2 was less numerous, but *A. lentulus* PWCAL1 was not annotated in mdr2 and mdr3. PDR subfamily classification showed that *A. lentulus* PWCAL1 was mainly annotated to cdr1 and AtrB, while the other three strains were mainly annotated to AtrF and AtrB without cdr1 and cdr2. The above results suggest that the distribution of different subfamilies of ABC proteins is also different between different *A. lentulus* strains.

In this study, the predicted ABC transporters provide some clues to investigate the resistance of clinically isolated azole resistant bacteria, however, many ABC subfamily proteins are remain unexplored. Our analysis can only predict their putative function. Therefore, based on ABC transporter family resistance gene prediction analysis, we suggest that the mechanism by which *A. lentulus* develops resistance to multiple azoles may be partially mediated by its own efflux system overexpression mechanism. We speculate that *A. lentulus* is a relatively new pathogen compared with *A. fumigatus*, with a lower infection rate than *A. fumigatus* and less external stimulation, so the gene sequence is relatively stable and the probability of genetic variation during evolution is rare, so the azole resistance mechanism caused by *A. lentulus* may be mainly based on the overexpression of its own efflux pump gene rather than CYP51 gene mutation, perhaps with the widespread use and overexposure of azoles, it may lead to the development of acquired resistance in *A. lentulus*.

## 4. Conclusion

In the present study, we obtained the fine mapping data of the whole genome sequence of *A. lentulus*, which provided detailed genomic information for a comprehensive understanding of the species at the molecular level, while the whole genome data of *A. lentulus* would also enrich the *A. fumigatus* complex species database and provide an important reference for the pathogenic control of *A. fumigatus* complex species. Based on the F46Y/N248T locus mutation combination identified by whole genome sequencing, we suggest that the CYP51A gene mutation in *A. lentulus* may be partially responsible for resistance to azoles. According to the predicted resistance gene analysis of ABC transporter family, we suggest that the mechanism of resistance to multiple azoles in *A. lentulus* is at least partially mediated by non-CYP51A gene resistance mechanisms.

## Author Contributions

All authors made a significant contribution to the work reported, whether that is in the conception, study design, execution, acquisition of data, analysis and interpretation, or in all these areas; took part in drafting, revising or critically reviewing the article; gave final approval of the version to be published; have agreed on the journal to which the article has been submitted; and agree to be accountable for all aspects of the work.

## Acknowledgements

This study was supported by grants from the Xinjiang Nature Science Foundation of China (2021D01E30) . We would like to thank all participants who participated in this study.

